# Tuning into the city soundscape: Optimizing Convolutional Neural Networks for avian acoustic identification in the neotropics and evaluating their performance against established monitoring approaches

**DOI:** 10.64898/2026.05.10.724049

**Authors:** Melissa Ardila-Villamizar, Laurence H. DeClippele, Davide M. Dominoni

## Abstract

Convolutional Neural Networks (CNNs) have become increasingly prominent in biodiversity monitoring due to their strong performance in accurately detecting species from sound recordings, overcoming some limitations of traditional methods such as point-counts. Yet, their use in urban ecosystems remains limited, highlighting the need for frameworks that identify modelling strategies to optimize their performance in these complex soundscapes. Here, we evaluated how preprocessing and labelling strategies, detection thresholds, sample size, and architecture affect the performance of CNNs for bird identification in urban tropical ecosystems. We also assessed its potential by comparing CNN-derived biodiversity estimates with those from point-counts and acoustic indices. For this, we used one week of recordings collected along urbanization gradients in five Colombian Andes cities to develop 11 multiclass CNN models varying in spectral representation, labelling strategies, training data source and backbone architecture. The best-performing model, evaluated with F1-scores, combined Log-Mel spectrograms, multispecies labels, ecosystem-specific and XenoCanto recordings, a probability threshold of 0.3 and a ConvNeXt backbone with its performance generally improving with sample size. Although CNNs and point counts detected partially distinct assemblages, CNN-derived species richness was higher to that estimated from point-counts. Additionally, the Normalized Difference Soundscape Index (NDSI) was positively associated with richness, suggesting its potential as a biodiversity proxy in tropical urban soundscapes. Overall, by identifying effective modelling designs and monitoring strategies, our study advances the development of robust biodiversity assessment frameworks in urbanized ecosystems in the Neotropics whilst highlighting the potential of acoustic approaches for avian monitoring and providing methodological guidance for future research and practical insights for wildlife monitoring and conservation.

## 1. Introduction

Although biodiversity declines are especially pronounced in urban ecosystems, monitoring efforts have traditionally focused on less disturbed environments (as suggested by Finnerty et al. 2025). Yet cities can support substantial biodiversity, particularly in the tropics, that contributes to ecosystem services such as air purification, climate regulation, and cultural enrichment (Kowarik et al., 2025). Moreover, certain groups such as birds can be particularly useful as bioindicators of ecosystem integrity and biodiversity, because of their sensitivity to environmental change and conspicuousness (Fraixedas et al., 2020). However, monitoring avian diversity in urban systems can be challenging, as their environmental heterogeneity and constant disturbance can affect both species presence and detectability (Van Heezik and Seddon, 2012). Highlighting the need of identifying monitoring methods that can reliably capture avian diversity under complex conditions especially in diverse but rapidly urbanizing tropical cities.

Bird monitoring typically relies on two methods, point-count surveys and passive acoustic monitoring (PAM). During point-counts, trained observers record the species they visually and acoustically identify within a defined time and distance (Venier et al., 2012). Thus, whilst point-counts are a widely applicable and valuable tool for generating ecological insights, especially for birds that vocalize infrequently, they can be time and resource-intensive, biased by observer experience and perception, and could underestimate diversity in structurally complex habitats (Farmer et al., 2012; Sliwinski et al., 2016). Contrastingly, PAM uses autonomous recording devices to capture soundscapes, thereby enabling species identification and, consequently, biodiversity assessment (Penar et al., 2020). Although PAM offers a continuous, non-invasive approach that can reduce observer bias and field effort, it generates large datasets and may be less effective for less vocal species (reviewed in Gibb et al. 2019). As point-counts and PAM differ in their strengths and limitations, evidence regarding the most effective approach for monitoring bird diversity remains mixed. Some studies have reported that point counts were able to detect more species than acoustic monitoring along elevational gradients in Australian rainforests (Leach et al., 2016), whilst others found that automated methods incorporating deep-learning tools such as BirdNET (Kahl et al., 2021) outperformed visual surveys in a temperate forest in Poland (Winiarska et al., 2025) and further analyses have shown that when differences in detection range and signal-to-noise ratio are accounted for, both methods can yield comparable outcomes in a metanalysis including different biomes (Darras et al., 2018) and in empirical studies in Afrotropical ecosystems (Joel et al., 2024; wa Maina and Njoroge, 2025). However, most comparative studies have been conducted in relatively less disturbed ecosystems, limiting their applicability to urban environments.

Monitoring urban ecosystems requires methods that can reliably detect birds under complex conditions. Passive acoustic monitoring (PAM) could be well-suited to this context because it enables standardized sampling across heterogeneous environments where point counts may be less effective due to sensory saturation (suggested by Arzberger et al. 2025). Moreover, the complexity of urban systems poses challenges not only for avian monitoring but also for the interpretation of its derived data that could be addressed, despite some limitations, through two methods commonly used in PAM-analysis, acoustic indices and deep learning models (Arzberger et al., 2025). Acoustic indices summarize temporal, spectral, and amplitude patterns in recordings (Bradfer-Lawrence et al., 2023). Due to their computational efficiency, more than 70 acoustic indices have been developed, including the Bioacoustic (BI), Acoustic Complexity (ACI), Acoustic Diversity (ADI), Acoustic Evenness (AEI), Normalized Difference Soundscape (NDSI), and Shannon entropy (H) indices (Alcocer et al., 2022). These indices capture different soundscape properties, including biophonic intensity (BI), temporal variability (ACI), the distribution and evenness of acoustic energy (ADI, AEI), the balance between biological and anthropogenic sounds (NDSI), and overall acoustic complexity (H) (summarized in Supplementary Table S1) (reviewed by Alcocer et al. 2022; Bradfer-Lawrence et al. 2025). However, by assuming a direct correlation between acoustic complexity and species diversity, acoustic indices may misinterpret soundscape elements such as anthropogenic noise as biological activity, producing unreliable diversity estimates in urban soundscapes (as suggested by Fairbrass et al. 2017). Consequently, alternative approaches such as deep learning models may provide a more accurate framework for bioacoustics monitoring in urban ecosystems.

Deep learning models, particularly Convolutional Neural Networks (CNNs), are a class of multilayer neural networks designed for classification tasks that have become increasingly prominent in biodiversity monitoring due to their strong performance (e.g., Ruff et al. 2020; Jalal et al. 2020; Kobayashi et al. 2021; Xie et al. 2022). Moreover, unlike acoustic indices, which provide indirect proxies of biodiversity, deep learning approaches can specifically detect species, allowing estimation of both diversity metrics and community composition (Hu et al., 2025). Recently, architectures originally developed for image classification such as VGG, ResNet and ConvNeXt have been applied to ecological monitoring through transfer learning, including a ResNet-derived model for identifying approximately 6,000 species (i.e., BirdNET. Kahl et al. 2021), and the incorporation of ConvNeXt and VGG16 architectures for the identification of nearly 10,000 and over 300 bird species respectively (Heinrich et al., 2025; Madake et al., 2023). However, performance can vary among architectures, ResNet tends to outperform VGG-architectures as its residual connections enable deeper learning, whereas comparisons with ConvNeXt remain limited despite its emergence as a high-performing architecture (He et al., 2016; Tang, 2025). Performance can also vary geographically as relevant acoustic features can differ among ecosystems (Ghani et al., 2025). Indeed, a recent global assessment reported that BirdNET achieved higher accuracy in temperate regions but lower predictive ability in Central and South America, Africa, and Asia (Funosas et al., 2026). Together, these differences suggest that pretrained models may not generalize well to complex or underrepresented ecosystems such as the urban tropics, indicating that locally trained CNNs may improve species detection there.

Given that relatively few CNNs have been developed to detect species in tropical urbanization gradients (but see Duta and Behdad 2023), identifying the factors that could improve their performance can be a first, crucial step for advancing their implementation there (as suggested by Dias et al. 2025). First, multiple preprocessing and labelling choices such as buffer zones around labels, class weighting, spectral transformation, data-augmentation using public libraries and the inclusion of multispecies and absence labels can influence CNN performance as they can affect detection accuracy, yet their effects remain poorly understood (summarized in Table 1). Moreover, additional factors such as class sample size and probability detection thresholds can also influence CNN performance (Benkendorf and Hawkins, 2020; Dias et al., 2025; Hestness et al., 2017). Larger sample sizes generally improve model performance by increasing the representation of signal variability (Benkendorf and Hawkins, 2020), however, performance may plateau once additional samples become redundant or noisy (Hestness et al., 2017). Similarly, CNN performance can be highly sensitive to the selection of the probability detection threshold, as this choice directly mediates the trade-off between precision and recall, influencing rates of false positives and missed detections (Knight et al., 2017; Stowell et al., 2019a). Therefore, evaluating the influence of architectures, probability thresholds, sample size, and preprocessing and labelling strategies on CNN performance allows to identify the most reliable approaches for species detection, strengthening CNN-based bird monitoring.

**Table 1.** Advantages and consequences of incorporating five model design choices to improve the performance of Convolutional Neural Network in species identification. These approaches were used to develop eight CNNs for bird species identification (see *Methods* section for more details about them).

| Approach | Advantages | Consequences |
| --- | --- | --- |
| 1. Adding buffer zones | Can soften label boundaries, making the model more flexible to identify calls under different acoustic scenarios (adapted from Lostanlen et al. 2019) | Can reduce label precision by including noninformative segments (Mu et al., 2021). |
| 2. Adding class weighting | Can improve model ability to detect rare species (Buda et al., 2018). | Can increase the detection of false positives (Buda et al., 2018) |
| 3. Using Log-Mel spectrograms | Highlights the low-mid frequencies of recordings where many bird vocalizations occur (LeBien et al., 2020) | Can be sensitive to background noise which could be overcome with PCEN spectrograms that intensify bird calls (Lostanlen et al., 2019b) |
| 4. Data augmentation with single-species labels | Providing high quality inputs from external libraries can highlight relevant acoustic features (Denton et al., 2021) | Might limit species detection in real acoustic conditions (Stowell et al. 2019) |
| 5. Adding absence data and multispecies labels | Can allow models to learn to discern between bird calls and other soundscape elements and facilitates species identification within overlapping vocalizations (Stowell et al., 2019a). | Including presence-data focus the model on learning specific features of targeted species, improving detection sensitivity (LeBien et al., 2020) |

This study aimed to 1) identify the preprocessing, labelling, threshold, sample size, and architectural choices that improve CNN performance for bird identification in tropical urbanization gradients, and 2) evaluate the potential of the best CNN to estimate biodiversity in challenging soundscapes by comparing its performance with a widely used CNN for bird identification, BirdNET (Kahl et al., 2021), and two commonly used approaches within the biodiversity monitoring framework, point-counts and acoustic indices. Specifically, we compared CNN-derived species richness with richness obtained from point counts and tested whether acoustic indices captured biodiversity patterns similar to those detected by the CNN, while accounting for interactions with impervious surface extent as a proxy for urbanization. For this, we conducted visual surveys and collected one week of acoustic recordings at 30 sites across urbanization gradients in five cities in the Colombian Andes. Then, we calculated six acoustic indices commonly used in urban ecosystems (NDSI, ACI, H, ADI, AEI, and BI) and developed 11 CNN models using the recordings. Eight models tested different preprocessing and labelling strategies, including spectral transformation, data augmentation, absence and multispecies labels, buffer zones, and class weighting. Then, the best configuration, identified using the F1-score, was tested across three architectures VGG16, ResNet, and ConvNeXt to select the optimal model. Finally, the top-performing CNN was compared with BirdNET, acoustic indices, and point-counts to assess its potential for biodiversity monitoring.

We hypothesized that CNN performance would improve with PCEN spectrograms, species weighting and the inclusion of multispecies and absence labels, as these approaches could address the strong class imbalance, overlapping vocalizations, and anthropogenic noise in our study system (adapted from Stowell et al. 2019). We also predicted that performance would increase with sample size before plateauing and, that lower detection thresholds (e.g., 0.1) would enhance true positive detection due to vocalizations being often partially masked by noise (Knight et al., 2017; Stowell et al., 2019a). Moreover, we expected locally trained CNNs to outperform pretrained architectures and BirdNET as these models are typically optimized on ImageNet or acoustic datasets from different ecological contexts, potentially limiting their performance in our study system (adapted from Ghani et al. 2025). Finally, given that the environmental complexity of urban ecosystems may hinder visual detection and inflate acoustic index values (Fairbrass et al., 2017), we predicted that our CNN would detect more species than point counts, that acoustic indices would not be related to CNN-derived richness, and that impervious surface extent would affect both point-count and CNN-derived richness by reducing species detection. Overall, by identifying effective modelling designs and monitoring strategies, our study advances the development of robust biodiversity assessment frameworks in complex urban environments in the Neotropics whilst also providing methodological guidance for future research and practical insights for wildlife monitoring and conservation in this diverse yet rapidly urbanizing landscapes.

## 2. Materials and Methods

### 2.1 Study area

We conducted point-count surveys and collected acoustic data in five cities across the Colombian Andes selected based on accessibility and to ensure broad representation of this region (Fig. 1). This data collection allowed us to gather the dataset used to build our Convolutional Neural Network models (CNNs), calculate acoustic indices and compare these approaches to evaluate the potential of the best-performing CNN.

**Fig. 1.**
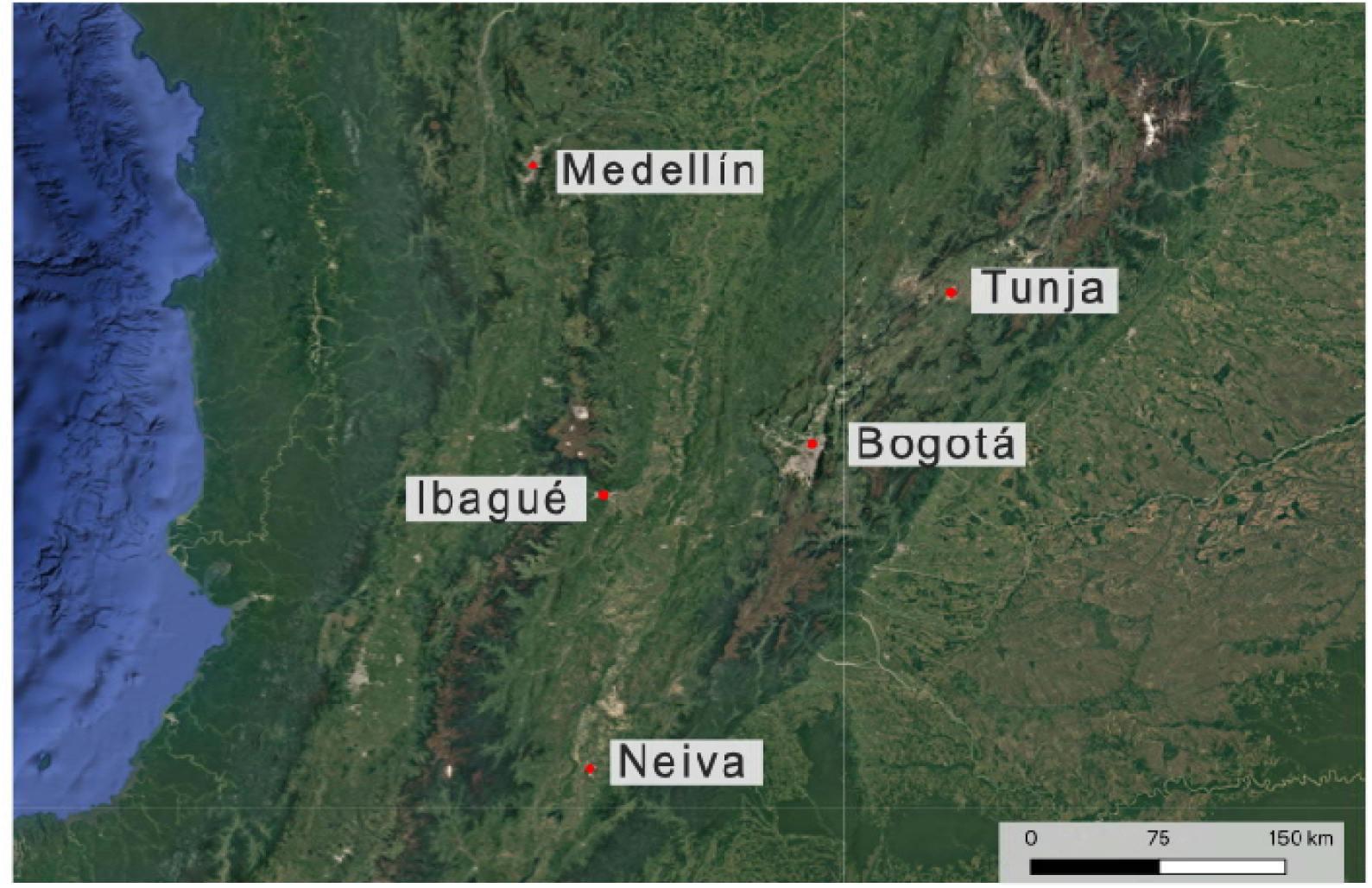
Map highlighting the five Colombian Andes cities selected for this study. Cities (Bogotá, Medellín, Ibague, Neiva and Tunja) are represented by a red dot. This image was recovered from Esri World Imagery.

In each city, we selected six sites (30 in total, Supplementary Fig. S1) to represent a local urbanization gradient. Urbanization gradients consisted of sites within the cities varying in impervious surface extension and patterns of human use, with some allowing recreational activities and others restricting them (Indicated in Supplementary Table S2). Thus, these urbanization gradients allowed us to evaluate the performance of the different monitoring approaches across varying levels of urbanization and human-induced disturbances. The impervious surface for each study site was calculated by generating a 500-meter buffer at each location to then use the Zonal Statistics tool in QGIS (QGIS Development Team, 2018) on the 2024 Global Artificial Impervious Area (GAIA) raster covering the study region (Gong et al., 2020) (Supplementary Table S2).

### 2.2 Acoustic and visual data collection

We deployed three automated recording units (AudioMoth v 5.2. Open Acoustic Devices 2025) that recorded continuously for a week within January and March 2025 at each study site, aiming to have an optimal representation of the local avian assemblages. The devices were deployed two meters above ground and located at least 80 meters apart from each other to minimize acoustic interference (following John Young et al., 2021).

Additionally, visual surveys (i.e., point counts) were performed by MAV along a 20x50 meter transect from each Audiomoth location by recording all the species seen or heard during a 10-minute period within birds’ peak activity hours (07:00–10:00 AM) (following Mitchell et al., 2020). Each transect was surveyed twice on different days.

### 2.3 Development of CNN models for bird-species identification in urban ecosystems

We trained, validated, and evaluated 11 Convolutional Neural Network (CNN) models to identify avian species across Neotropical urbanization gradients. Eight models were used to systematically evaluate how preprocessing and labelling choices, probability thresholds, and species sample size affect CNN performance, whereas the remaining three were used to compare alternative architectures and identify the best-performing model for this case study.

To determine the optimal model design, we implemented a stepwise evaluation framework (Fig. 2) addressing common challenges in passive acoustic monitoring, including background noise, class imbalance, and annotation uncertainty (adopted from Stowell et al. 2019). Across five stages, we sequentially assessed the influence of spectral representation (Log-Mel vs. PCEN), data augmentation, labelling strategies (including presence-absence data and multispecies labels), class weighting, and temporal buffering on CNN performance. This sequence followed the typical development pipeline of a CNN, progressing from fundamental input and data representation to progressively finer labelling adjustments, allowing us to isolate the effect of each modelling decision while minimizing potential confounding effects.

**Fig. 2.**
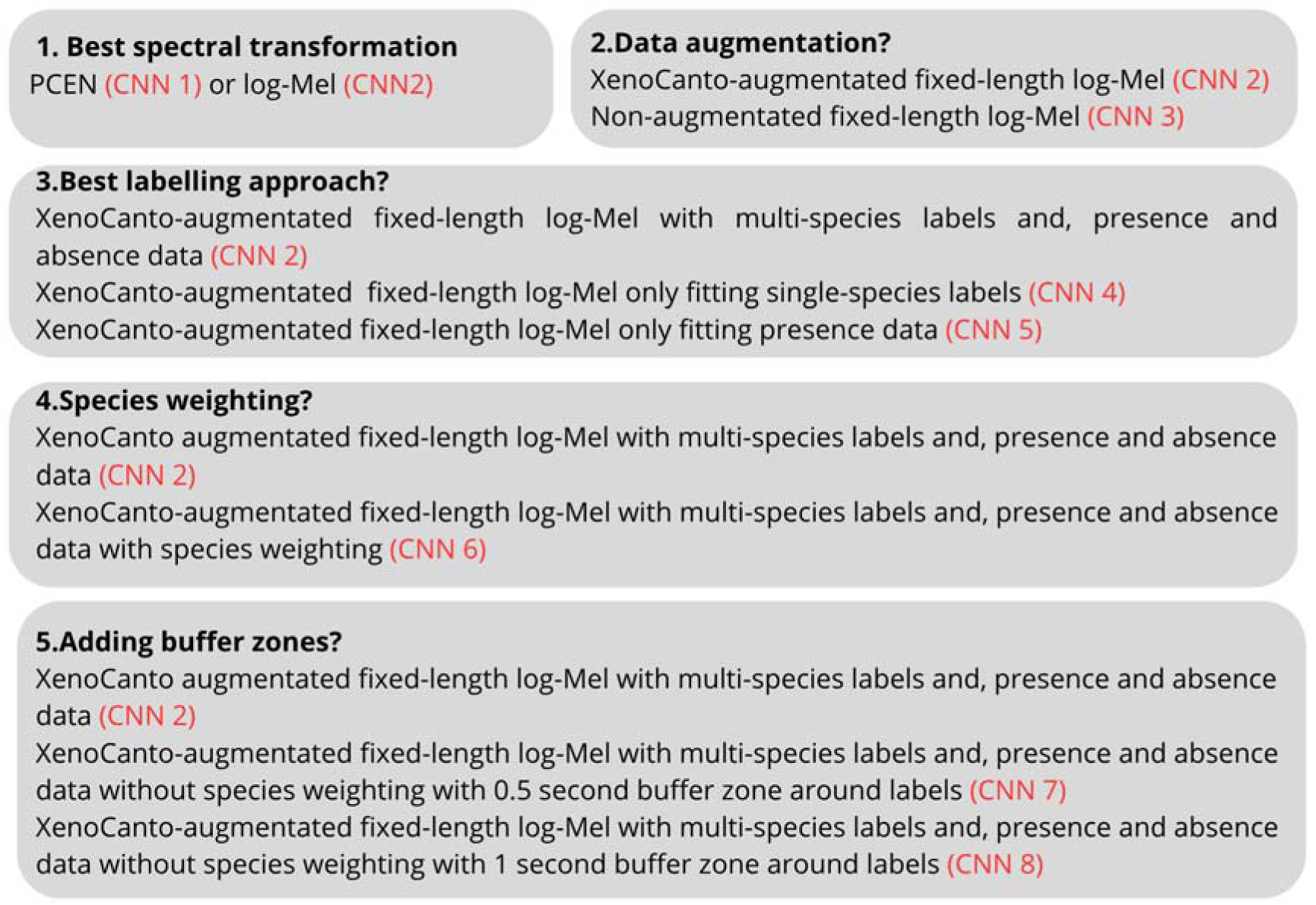
Scheme illustrating the stepwise evaluation framework used to assess model design choices for a bird-species identification CNN developed for tropical urban-rural gradients. Eight distinct model configurations were evaluated to identify the design yielding the best performance.

Following this evaluation framework, we first compared spectral representations by training two CNNs using PCEN (CNN 1) and Log-Mel spectrograms (CNN 2). Both models were trained with a combination of field recordings and XenoCanto data (Xeno-canto Foundation, 2025) and included presence and absence labels representing non-avian sounds to reflect common monitoring scenarios. Second, we assessed the effect of data augmentation by comparing the best-performing model (CNN 2) with one trained exclusively on field recordings (CNN 3). Third, we evaluated labelling strategies by comparing a model trained with multispecies and presence-absence labels (CNN 2) with models using single-species labels (CNN 4) and presence-only data (CNN 5). Fourth, we tested whether incorporating class weighting improved performance by comparing a model previously trained without species-specific weighting in the loss function (CNN 2) with one addressing it (CNN 6). Multiple upper limits for positive class weighting (5, 8, and 10) were evaluated (following Razali et al. 2025), with a weight of 8 providing the best balance between sensitivity to underrepresented species and training stability. Lastly, we assessed the effect of temporal buffering by adding 0.5 (CNN 7) and 1 second silence buffers (CNN 8) around labelled vocalizations and compared them with no buffering (CNN 2), following biologically relevant call durations (Priyadarshani et al., 2018). Model development involved four stages: data labelling, data augmentation, spectrogram generation, and model training and evaluation.

#### 2.3.1 Data labelling

First, we generated the dataset to train, validate and test the performance of all the developed CNN models by manually labelling segments of randomly selected five-minute fragments from each recording hour of one day at each AudioMoth recording location (90 nodes), to indicate the presence or absence of an identified avian species using RavenPro 1.6 (K. Lisa Yang Center for Conservation Bioacoustics at the Cornell Lab of Ornithology., 2025). To avoid errors during the labelling phase, the identity of the birds present in the segments was always confirmed by comparing its spectrogram with others available in XenoCanto. In total, our training dataset consisted of 10.800 minutes or 180 hours of 17,979 labelled segments from 147 species of 16 orders, 35 families and 116 genera (species and number of labels per species are summarized in Supplementary Table S3).

#### 2.3.2 Data augmentation

We retrieved vocalizations for each of the 147 species identified during the labelling phase from the widely used community bird-sound library XenoCanto (Xeno-canto Foundation, 2025) for the model assessing the effect of including external labels in CNN performance. For this, we retrieve audio files through the XenoCanto public API (Xeno-canto Foundation, 2025) using R v 4.5 (R Core Team, 2021). Recordings were collected from Colombia and four neighboring South American countries: Brazil, Ecuador, Peru and Venezuela, to capture natural regional variation in vocalizations while avoiding the inclusion of distant dialects that could hinder model learning. The search iterated through these countries until reaching 1000 samples per species. The country, date, time, XenoCanto ID, URL and author of each file was recorded in a metadata base to avoid duplication and keep relevant recording details.

#### 2.3.3 Spectrogram computation

After obtaining the labelled training dataset, we converted recordings to either Log-Mel or Per-Channel Energy Normalized (PCEN) spectrograms as one of our models aimed to assess the best spectral transformation. For this, we first fragmented recordings into fixed-length five-second segments and normalized them using the global mean and standard deviation of the time-frequency bins computed across the entire dataset aiming to achieve scale consistency across samples and stabilize model training (following LeBien et al. 2020). Then, for the dataset following the Log-Mel transformation, we resampled all the recordings to 48 kHz and computed spectrograms using a 2048-point fast Fourier transform, a hop length of 480 samples, and 128 Mel frequency bands (adapted from Baowaly et al. 2024). PCEN spectrograms were first transformed to the Log-Mel scale using the same spectral parameter (frequency, hop length and fast Fourier transformation) and then, the instantaneous power of each band was normalized for each band using an exponential moving average for adaptative gain control followed by dynamic range compression (adapted from Michaud et al. 2025). All spectrograms were computed using the Python v 3.12 modules numpy v 2.3.5, soundfile v 0.13.1, resample_poly v 1.16.2 and librosa v 0.11.0 (Bechtold, 2020; McFee et al., 2015).

#### 2.3.4 Model training and testing

We used the spectrograms generated in the previous step as the input of eight Convolutional Neural Networks (CNNs) that enable the identification of preprocessing and labelling strategies that yielded the best performance in bird-species identification. All models used the same architecture to allow controlled comparisons among them.

CNN architecture was adapted from (Duta and Behdad, 2023; Salamon and Bello, 2017; Stowell et al., 2019b) and comprised three convolutional layers followed by global pooling and a fully connected output layer. Each convolutional block consisted of a two-dimensional convolution followed by a rectified linear unit (ReLU) nonlinearity. The first convolutional layer used 24 filter with a kernel size of 5 x 5, stride 2 and padding of 2. The second and third convolutional layers used 48 and 96 filters, respectively, each with a kernel size of 3 × 3, stride 2 and padding of 1. The use of stride convolutions progressively reduced the spatial resolution of the feature maps while increasing the number of feature channels, allowing to progressively extract more abstract acoustic features while remaining computationally efficient (Gunawan et al., 2021). Following the convolutional layers, feature maps were compressed using adaptive average pooling, resulting in a fixed-length feature vector of dimension 96, independent of the temporal duration of the input spectrogram. Feature maps were then fed into a fully connected linear layer that produced one output neuron per targeted species. Then, a sigmoid activation function was applied to each output neuron to produce probabilities between 0 and 1, representing the presence (1) or absence (0) of each species. Furthermore, the model was trained under a multi-label classification setting in all the models expect from the one only including single-species labels, allowing it to detect multiple bird species within a single recording, thereby making the model robust to real life acoustic conditions (as suggested by Knight et al. 2017).

Training, validation and test datasets were built under a per-species stratified split that included 70% of the data for each species for training, 20% for validation and 10% for testing, ensuring that all species were represented across splits (adapted from Stowell et al. 2019). Lastly, model optimization was conducted using the Adam optimizer with a learning rate of 1 × 10⁻³ including binary cross-entropy loss as the objective function (Kingma and Ba, 2017). Each model was trained three times with independent random initializations, using seeds 7, 42, and 1337 to account for stochastic variability in neural network training (Bouthillier et al., 2021). Lastly, model performance was evaluated by applying a sigmoid function to the network outputs to obtain species-level detection probabilities for the test set (following O’Shea and Nash 2015). Performance metrics were computed under three probability thresholds (0.1, 0.3, and 0.5) to assess the ability of the model to detect species (recall) and its accuracy when making these inferences (precision) (O’Shea and Nash, 2015). Then, overall performance was summarized using the micro-averaged F1-score, calculated by aggregating true positives, false positives and false negatives across all classes, reflecting CNN performance.

Models were developed using Python v 3.12 models using the packages PyTorch, Torchvision, NumPy, SoundFile, SciPy, and Librosa (Bechtold, 2020; Imambi et al., 2021; McFee et al., 2015; Virtanen et al., 2020).

#### 2.3.5 Identifying the model architecture that enhances CNN performance

Once the optimal model design was identified, we evaluated its performance across three pretrained CNN backbones: VGG16 (Simonyan and Zisserman, 2014), ResNet (He et al., 2016) and ConvNeXt (Liu et al., 2022), using transfer learning to determine the most effective architecture. These backbones were selected due to their widespread adoption and demonstrated effectiveness in spectrogram-based classification tasks (Dias et al., 2025; Gunawan et al., 2021; LeBien et al., 2020; Madake et al., 2023). For this comparison, we adopted the modelling design and dataset of the best-performing configuration (CNN 2), which used Log-Mel spectrograms, included both presence and absence data with multispecies labels, and excluded data augmentation, species weighting, and temporal buffer zones. Then, the original CNN architecture was replaced with each pretrained backbone, while all other design components were held constant to ensure a fair comparison. In each model, the original classification layers of the pretrained networks were removed and replaced with a task-specific species classification head (following Heinrich et al. 2025). The pretrained backbones acted as shared feature extractors, while newly initialized fully connected layers were trained to predict species presence. Because these architectures were originally designed for three-channel RGB images, the first convolutional layer was adapted to accept single-channel spectrogram inputs by averaging pretrained weights across input channels, enabling reuse of learned spatial filters for acoustic representations (Heinrich et al., 2025).

Model performance was evaluated using the same criteria described in the *Model training and testing* section, with a fixed probability threshold of 0.3, which yielded the best overall performance and allowed for direct comparison between the model trained from scratch and those employing transfer learning.

### 2.4 Comparison of the best performing CNN and BirdNET

To contrast our model against a widely used automated species identifier, we compared predictions from our best-performing model: CNN 2 implementing a ConvNeXt backbone (mean F1-score= 0.813±0.268, see Supplementary Table S3 for the F1-scores of each of the included species), with those generated by BirdNET using the BirdNET-Analyzer open-source tool (Kahl et al., 2021). To conduct this comparison and establish reference expert-validated annotations, we compiled a one-hour audio recording that was not used during model training or testing consisting of randomly selected two-minute segments from each study site. Then, to establish expert-validated annotations, species present in this compilation were manually labelled by MAV and an external ornithologist with regional expertise to provide an external reference (Dr. Jaime Andres Garizabal Carmona). Ground-truth annotations were established using a consensus approach, where a species was considered present only when identified by both the expert and the authors.

After getting the reference data, we ran each model to generate species-level predictions within the one-hour audio compilation at a probability threshold of 0.3 as it yielded the highest overall performance for our model and corresponds to the default detection threshold used by BirdNET (Kahl et al., 2021). Then, the predictions made by each model were compared against the expert-validated reference annotations by quantifying the total number of inferences, true positives, and false positives made by each model and thus, evaluate their overall performance through precision, recall, and both micro and macro-averaged F1-scores, computed relative to the reference annotations. Due to the limited number of comparative samples, no statistical significance testing was performed when comparing expert annotations to BirdNET and CNN predictions.

### 2.5 Quantification of acoustic indices

To compare biodiversity estimates derived from acoustic data with those obtained from our best-performing CNN model and point counts, we calculated six commonly used acoustic indices relevant for urban biodiversity monitoring: the Normalized Difference Soundscape Index (NDSI), Acoustic Complexity Index (ACI), Shannon entropy (H), Acoustic Diversity Index (ADI), Acoustic Evenness Index (AEI), and Bioacoustic Index (BI) (Arzberger et al., 2025; Fairbrass et al., 2017; Santos et al., 2024)

To quantify each index, we first segmented one week of recordings from each study site using a sliding-window approach (60 s windows with 10 s overlap) to capture temporal variation in soundscape activity where each section was resampled to a standardized sampling rate of 48 kHz prior to feature extraction. We then transformed each segment into a power spectrogram using a short-time Fourier transform (STFT), from which acoustic indices were calculated (following Bradfer-Lawrence et al. 2019). Lastly, we averaged index values across segments to generate mean values per site that were used as biodiversity proxies in subsequent analyses. Indices were computed using the Python v 3.12 modules numpy (Bechtold, 2020), scipy (Virtanen et al., 2020), librosa (McFee et al., 2015), and soundfile (Bechtold, 2020).

### 2.6 Data analysis

#### 2.6.1 Identification of the best CNN model

Initially, we fitted two generalized models to identify the CNN configuration with the best preprocessing, labelling strategies, and architecture. First, we fitted a Generalized Linear Mixed Model using Template Model Builder with a beta-regression distribution to examine how different modelling choices influenced CNN performance using the glmmTMB package v 1.1.13 (Bolker, 2019). The model fitted F1-score as the response variable and included model type (Log-Mel, PCEN, single-species and presence labels, including species-weighting and incorporating a 0.5 or 1 second buffer zone), random seed, probability threshold used for evaluation and the log-transformed number of samples per species as predictors. An interaction between model type and sample size was also included because sample size varied systematically across model types. Sample size was log-transformed to reduce skewness and account for diminishing returns in model performance with increasing data availability, thereby enabling a more interpretable relationship with performance (as suggested by Knight et al. 2017). The dataset used in this analysis comprised 10,584 F1-score per-species observations across all evaluated models, including three random seeds and three probability thresholds.

Because the relationship between sample size and F1-score remained nonlinear even after log-transformation, we additionally fitted a Generalized Additive Mixed Model (GAMM) to account for potential nonlinear effects of sample size using the gamm function of the mgcv package v 1.9-4 (Wood and Wood, 2015). The GAMM included the same predictors as the initial GLMM but modelled log-transformed sample size using a smooth function (k = 10). The fit of both models was then compared using Akaike’s Information Criterion (AIC), which indicated that the GAMM provided a better fit to the data (ΔAIC ≈ 399). Consequently, the GAMM was retained as the primary model for inference, and its estimates are reported in the *Results* section. Lastly, we performed post hoc pairwise comparisons among model configurations to evaluate differences in performance between them using the emmeans package v2.0.1 (Lenth et al., 2018).

For the second model aiming to assess the influence of four different architectures in the CNN performance fitting F1-score, we only included the F1-score observations from the best model design assessed in the previous step and compared them with the F1-score observations across the other three evaluated pre-trained architectures including three random seeds and one probability threshold (0.3) as it yielded the best performance. This model fitted F1-score as a response variable and architecture (own, ResNet, VGG16 or ConvNeXt) as a predictor and comprised 1323 observations.

All models fitted seed and species as a random effect and their overall performance regarding model convergence, collinearity, residual diagnostics, and potential heteroscedasticity was assessed by checking residuals patterns and using the performance package v 0.15.3 (Lüdecke et al., 2021). Data analysis was performed using R v 4.5 (R Core Team, 2021).

#### 2.6.2 Comparing biodiversity estimates across monitoring methods

To evaluate the potential of our best-performing CNN model with alternative monitoring approaches, we compared species richness estimated using the CNN with species richness obtained from point-count surveys and biodiversity proxies from the acoustic indices. We focused on species richness rather than other diversity metrics because it allowed us to consistently compare across the three diversity approaches as acoustic indices primarily provide proxies for overall acoustic richness and do not allow reliable estimation of community composition or species identities (Giuliani et al., 2024). For this, we first estimated species richness at each study site as the number of unique species detected by applying the CNN to one week of recordings from each recording site to detect species (machine-learning richness) and using the species previously recorded during the point counts (point-count richness).

After estimating site-level species richness from CNN inferences and point-count surveys, we compared the two approaches using a Generalized Linear Model with a Poisson distribution using the glm function from base R that fitted richness as the response variable and method as a predictor along with an interaction between method and impervious surface extent to assess whether urbanization influenced richness estimates differently across approaches.

Then, to examine whether acoustic indices captured biodiversity patterns similar to those detected by the CNN, we fitted a Negative Binomial Generalized Linear Model (GLM), due to the overdispersion of the data, with machine-learning richness as the response variable and scaled acoustic indices: ACI, H, ADI, NDSI, and BI as predictors using the “glm.nb” function from the package MARSS v 7.3-65 (Holmes et al., 2012). This model comprehended a dataset of 30 observation representing the richness of each study site (N=30). As in previous steps, we evaluated model overall performance regarding collinearity, convergence, residual diagnostics, and potential heteroscedasticity by checking residuals patterns and using the performance package v 0.15.3 (Lüdecke et al., 2021). Here, we excluded AEI from the analysis due to its expected collinearity with ADI, as it is calculated as its inverse, and retained ADI because it is more commonly used as a proxy for biodiversity in bioacoustic studies (as suggested by Zhao et al. 2019). Data analysis was performed using R v 4.5 (R Core Team, 2021).

## 3. Results

In total, we developed eleven CNN models and conducted 180 point-count surveys. CNN models were developed to identify 147 species from 16 orders and 34 families in tropical urban-rural gradients (see Supplementary Table S3 for the species included), eight to identify the best model design and three to assess its optimal architecture. The mean performance, assessed through F1-score, was 0.621±0.14 for the best performing model (LogMel) and, 0.13±0.252 for the worst one (single data), with an average of 1160.301± 1001.976 samples per species across models (Table 2). Model performance varied with the probability threshold applied, with mean F1-scores of 0.353±0.253 at a threshold of 0.1, 0.358±0.267 at 0.3, and 0.323±0.268 at 0.5 in the dataset encompassing performance under different thresholds. Lastly, model performance varied across the evaluated architectures with VGG16 providing the worst performance (0.526±0.235) and ConvNext the best (0.813±0.268) (Table 2).

**Table 2.**
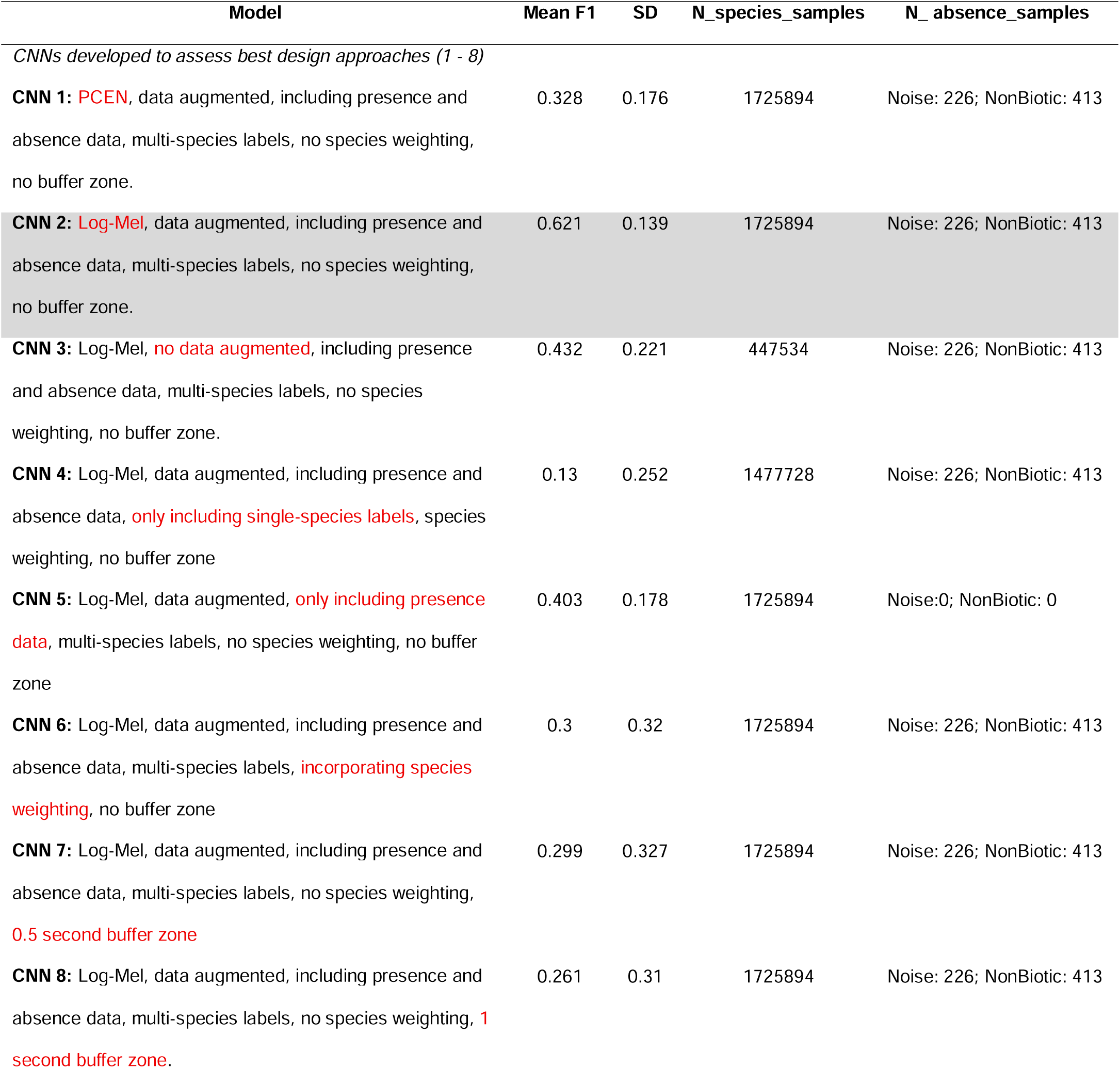
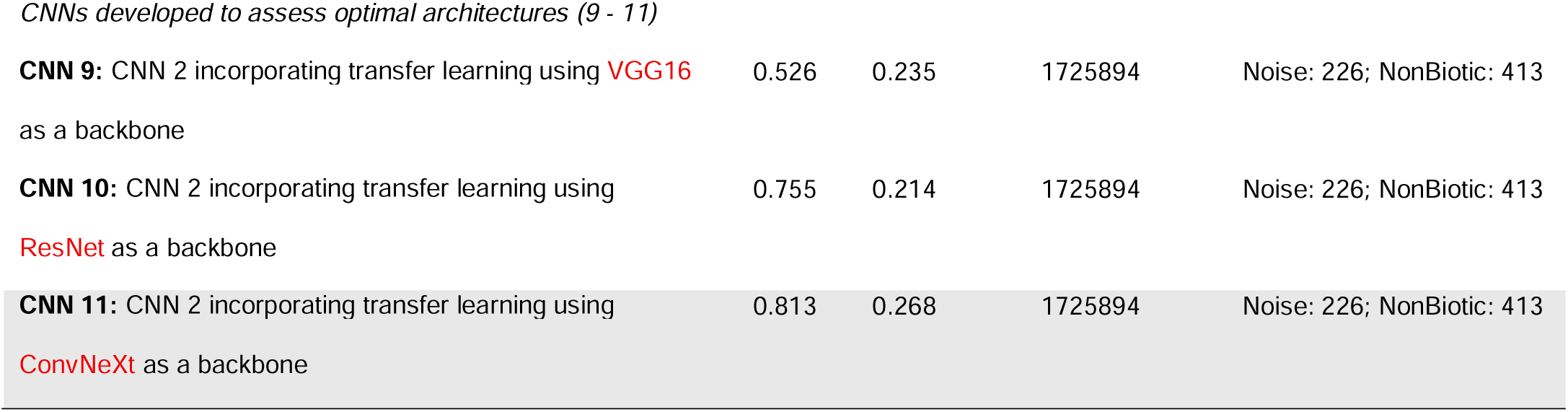
Summary of the number of species samples (n_species_samples), number of noise and non-biotic samples (N_absence_samples) and, mean and standard deviation of the performance (assessed as F1-score) of each of the 11 CNN models designed in this study. Model design choices and architectures incorporated in each model are highlighted in red, and the model with the best-performing design and architecture is shaded in grey.

| Model | Mean F1 | SD | N_species_samples | N_absence_samples |
| --- | --- | --- | --- | --- |
| *CNNs developed to assess best design approaches (1 - 8)* |  |  |  |  |
| **CNN 1:** PCEN, data augmented, including presence and absence data, multi-species labels, no species weighting, no buffer zone. | 0.328 | 0.176 | 1725894 | Noise: 226; NonBiotic: 413 |
| **CNN 2:** Log-Mel, data augmented, including presence and absence data, multi-species labels, no species weighting, no buffer zone. | 0.621 | 0.139 | 1725894 | Noise: 226; NonBiotic: 413 |
| **CNN 3:** Log-Mel, no data augmented, including presence and absence data, multi-species labels, no species weighting, no buffer zone. | 0.432 | 0.221 | 447534 | Noise: 226; NonBiotic: 413 |
| **CNN 4:** Log-Mel, data augmented, including presence and absence data, only including single-species labels, species weighting, no buffer zone | 0.13 | 0.252 | 1477728 | Noise: 226; NonBiotic: 413 |
| **CNN 5:** Log-Mel, data augmented, only including presence data, multi-species labels, no species weighting, no buffer zone | 0.403 | 0.178 | 1725894 | Noise:0; NonBiotic: 0 |
| **CNN 6:** Log-Mel, data augmented, including presence and absence data, multi-species labels, incorporating species weighting, no buffer zone | 0.3 | 0.32 | 1725894 | Noise: 226; NonBiotic: 413 |
| **CNN 7:** Log-Mel, data augmented, including presence and absence data, multi-species labels, no species weighting, 0.5 second buffer zone | 0.299 | 0.327 | 1725894 | Noise: 226; NonBiotic: 413 |
| **CNN 8:** Log-Mel, data augmented, including presence and absence data, multi-species labels, no species weighting, 1 second buffer zone. | 0.261 | 0.31 | 1725894 | Noise: 226; NonBiotic: 413 |
| CNNs developed to assess optimal architectures (9 - 11) |  |  |  |  |
| **CNN 9:** CNN 2 incorporating transfer learning using **VGG16**<br>as a backbone | 0.526 | 0.235 | 1725894 | Noise: 226; NonBiotic: 413 |
| **CNN 10:** CNN 2 incorporating transfer learning using<br>**ResNet** as a backbone | 0.755 | 0.214 | 1725894 | Noise: 226; NonBiotic: 413 |
| **CNN 11:** CNN 2 incorporating transfer learning using<br>**ConvNeXt** as a backbone | 0.813 | 0.268 | 1725894 | Noise: 226; NonBiotic: 413 |

On average, the best-performing CNN uniquely detected 103 species, with a mean species richness of 21.63 ± 9.256 species across study sites. In comparison, point-count surveys uniquely detected 84 species, with a mean richness of 14.23 ± 6.52 species. The mean values of the acoustic indices across sites were: NDSI −0.75 ± 0.24, ACI 800.79 ± 12.79, ADI 0.75 ± 0.09, AEI 0.51 ± 0.09, BI 198.01 ± 193.06, and H 0.88 ± 0.06.

### 3. 1 Identification of the best CNN model

We found that model performance, assessed using species-level F1-scores, was significantly influenced by model design, probability threshold and species sample size (Table 3, Supplementary Table S4). Among the evaluated designs, CNN 2 (referred as LogMel model) achieved the highest overall performance (Fig. 3), suggesting that using Log-Mel as a spectral transformation, not applying species weighting or buffer zones, using presence and absence, and multi-species labels that incorporated recordings collected both in the study system and from the external library XenoCanto improves the ability of CNN models to detect and correctly identify species in tropical urban-rural gradients.

**Fig. 3.**
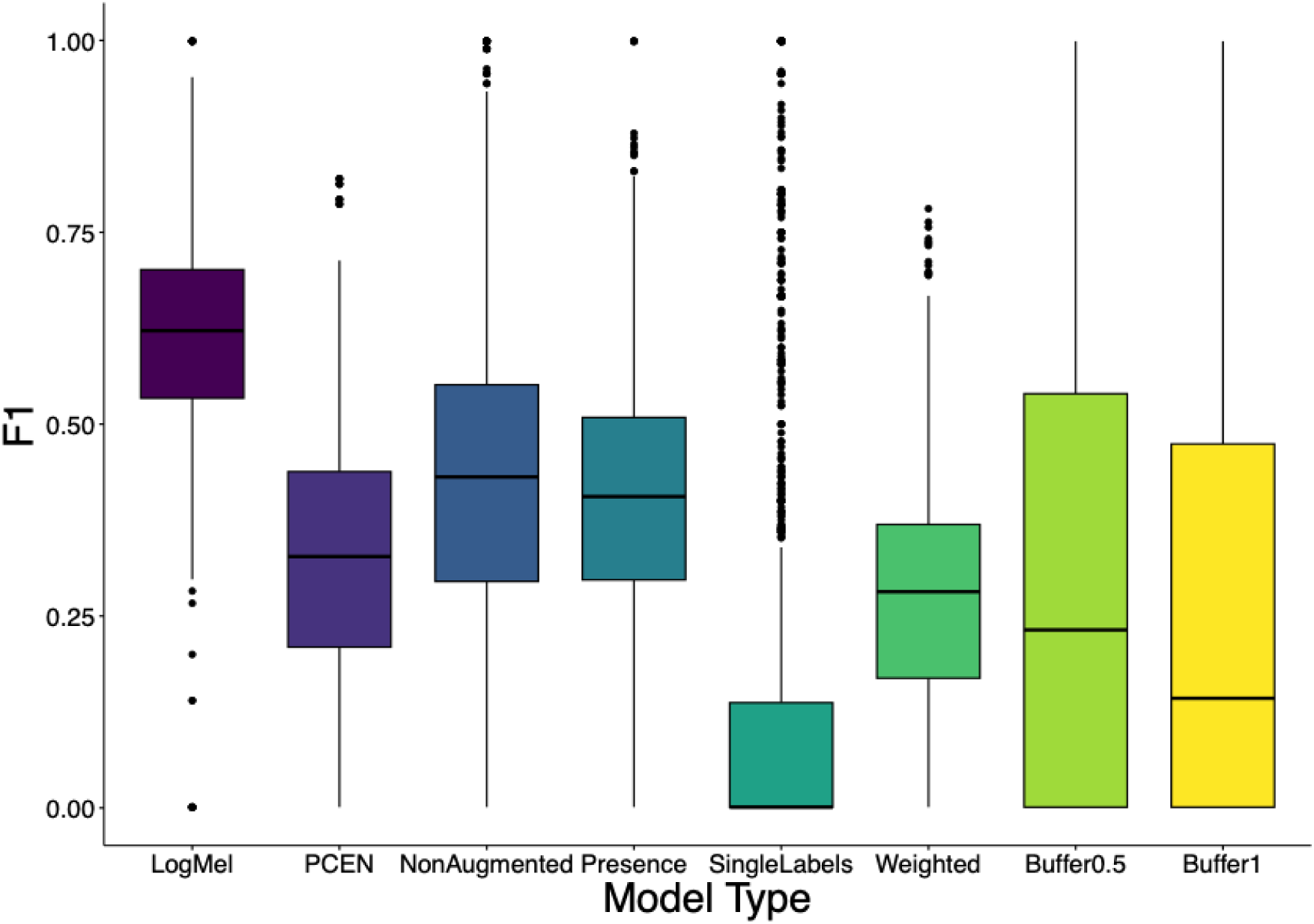
Differences in the performance (F1-score) of the eight CNN models developed to identify the best design. Each model is represented by a bar: LogMel (represented in dark purple, N=1323), PCEN (in light purple, N=1323), NonAugmented (in blue, N=1323), Presence (in turquoise, N=1323), SingleLabels (in dark green, N=1323), Weighted (in light green, N=1323), Buffer05 (in lime-green, N=1323) and Buffer1 (in yellow, N=1323). The plot shows the medians (black line inside the box) with boxes showing the lower and upper quartile, black vertical bars showing the full range of the data and dots outside the boxplot representing outliers. N= 10,584 corresponds to the total data included in the model. Pairwise comparison among model configurations can be find in Supplementary Table S3.

**Table 3.** Coefficients, confidence intervals and p-values from a Generalized Additive Mixed Model (GAMM) evaluating the influence of model design, threshold, log-transformed sample size and an interaction between model type and sample (fixed effects) on model performance (assessed as F1-score). The reference level for this model was the Buffer0.5 design. Bold values indicate statistical significance at α=0.05. N=10,584 corresponds to the total number of observations included in the regression model with 22 degrees of freedom. The variance explained by the fixed effect was R^2^ = 0.455. Model AIC was -16926.77.

| Predictors | Estimates | CI | P-value |
| --- | --- | --- | --- |
| Intercept | 0.17 | 0.09 – 0.31 | **<0.001** |
| **Buffer1** | **0.35** | **0.19 – 0.64** | **0.001** |
| **LogMel** | **45.18** | **24.95 – 81.79** | **<0.001** |
| **NonAugmented** | **14.07** | **8.09 – 24.47** | **<0.001** |
| **PCEN** | **11.58** | **6.41 – 20.92** | **<0.001** |
| **Presence** | **18.42** | **10.21 – 33.22** | **<0.001** |
| **Single** | **0.53** | **0.30 – 0.93** | **0.026** |
| **Weighted** | **14.21** | **7.86 – 25.67** | **<0.001** |
| **Threshold** | **0.58** | **0.52 – 0.65** | **<0.001** |
| **Log (Count)** | **1.09** | **1.00 – 1.19** | **0.039** |
| **Buffer1 × Log (Count)** | **1.14** | **1.05 – 1.24** | **0.003** |
| **LogMel × Log (Count)** | **0.74** | **0.68 – 0.81** | **<0.001** |
| **NonAugmented × Log (Count)** | **0.73** | **0.67 – 0.79** | **<0.001** |
| **PCEN × Log (Count)** | **0.78** | **0.71 – 0.84** | **<0.001** |
| **Presence × Log (Count)** | **0.76** | **0.70 – 0.82** | **<0.001** |
| Single × Log (Count) | 0.99 | 0.91 – 1.07 | 0.750 |
| **Weight × Log (Count)** | **0.74** | **0.68 – 0.81** | **<0.001** |
| Smooth term (species) |  |  | **<0.001** |
| Smooth term (seed) |  |  | **0.002** |

Furthermore, a probability threshold of 0.3 consistently yielded the best performance across model types, suggesting that in ecosystems where biotic signals may be weaker due to other soundscape components such as anthropogenic noise (e.g., urban environments), lowering the detection threshold can improve species identification (Fig. 4). However, this effect was relatively small (Fig. 4), indicating that model performance was only weakly sensitive to the probability detection threshold.

**Fig. 4.**
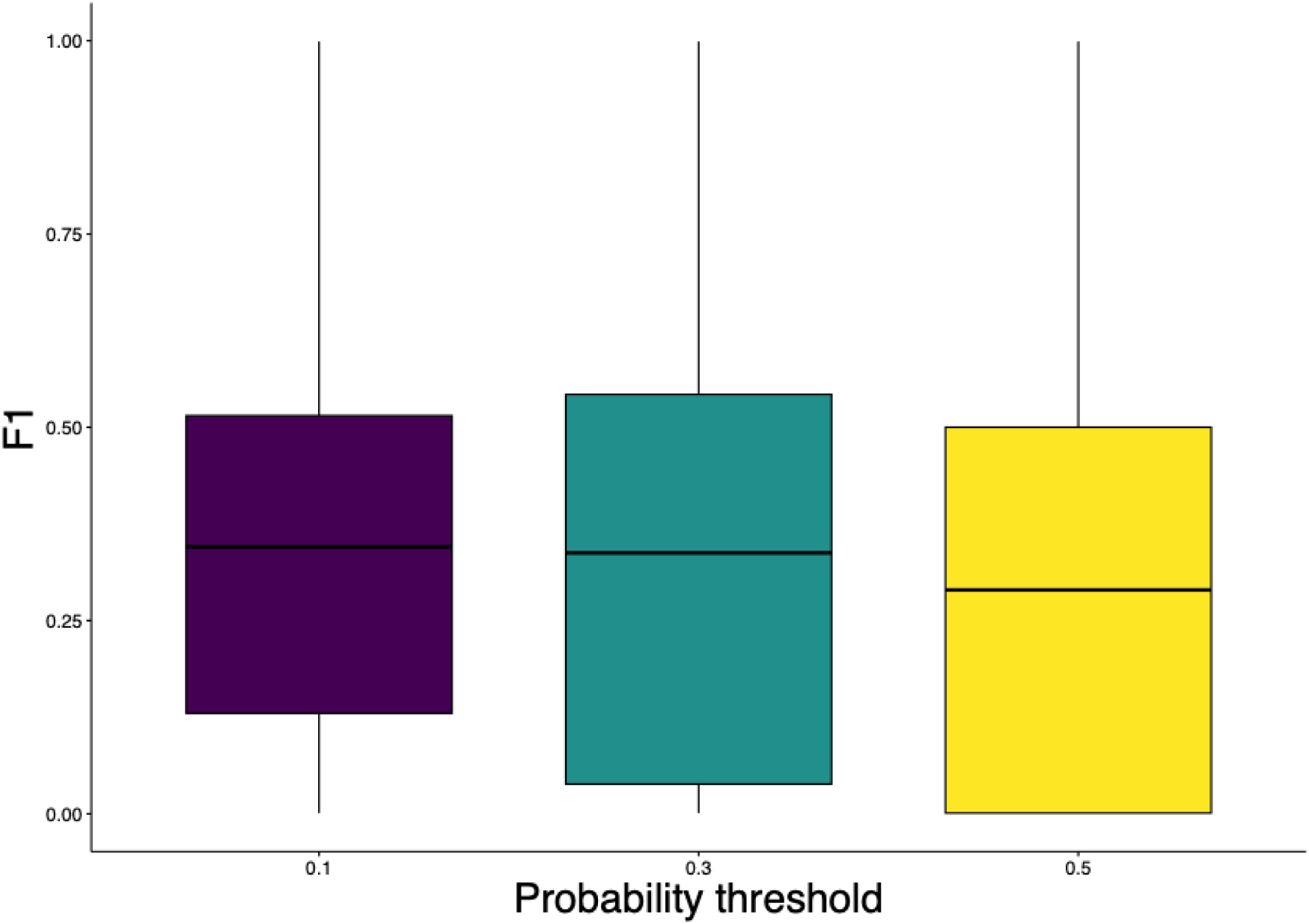
Differences of model performance (assessed as F1-score) across probability thresholds. Each probability is represented by a bar: 0.1 (represented in dark purple, N= 3528), 0.3 (in turquoise, N= 3528), and 0.5 (in yellow, N= 3528). The plot shows the medians (black line inside the box) with boxes showing the lower and upper quartile and black vertical bars showing the full range of the data. N=10,584 corresponds to the total data included in the model.

Moreover, species sample size exhibited a significant positive relationship with model performance (assessed as F1-score), suggesting that models trained with larger datasets achieved higher classification accuracy, although this improvement tended to plateau at higher sample sizes (Fig. 5, Table 3). However, the strength of this relationship differed among the evaluated model designs (Fig. 6, Table 3). Specifically, the performance of the model configurations LogMel, PCEN, NonAugmented, Presence and Weighted decreased with sample size whilst the performance of the models Buffer0.5 and Buffer1 increased as sample size increase. Finally, the interaction between sample size and the Single configuration showed a positive relationship but was not statistically significant. Overall, these results suggest that the benefits of increasing species sample size depend strongly on model design choices, with certain labelling strategies and temporal buffering approaches enabling more effective use of additional training data, while other configurations reach performance saturation more rapidly.

**Fig. 5.**
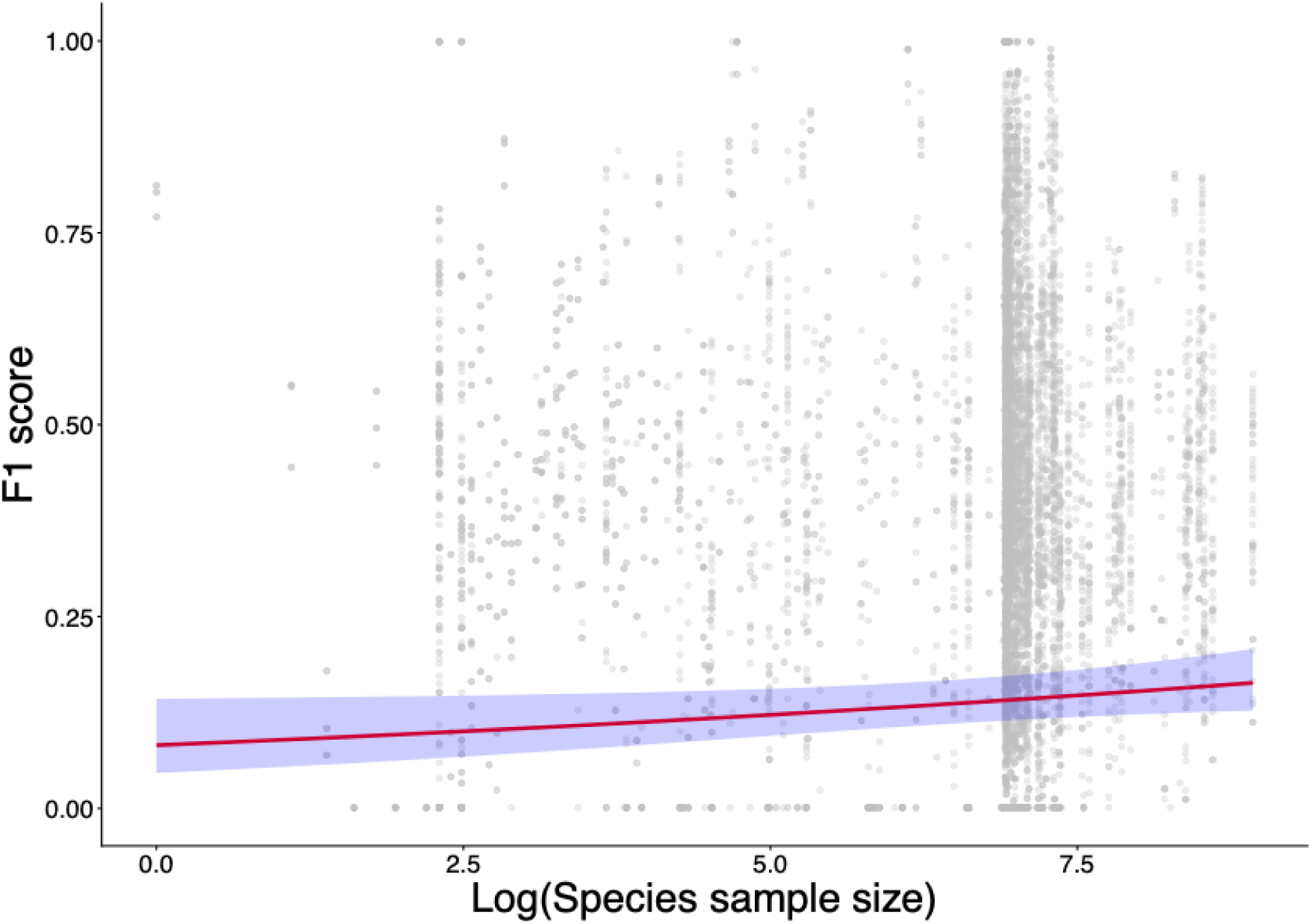
Relationship between log-transformed sample size and model performance (F1-score) (N = 10,584). The red line represents the predicted effect from the Generalized Additive Mixed Model, while the blue shaded area indicates the corresponding 95% confidence interval. Grey dots represent the raw observations (log-transformed sample size).

**Fig. 6.**
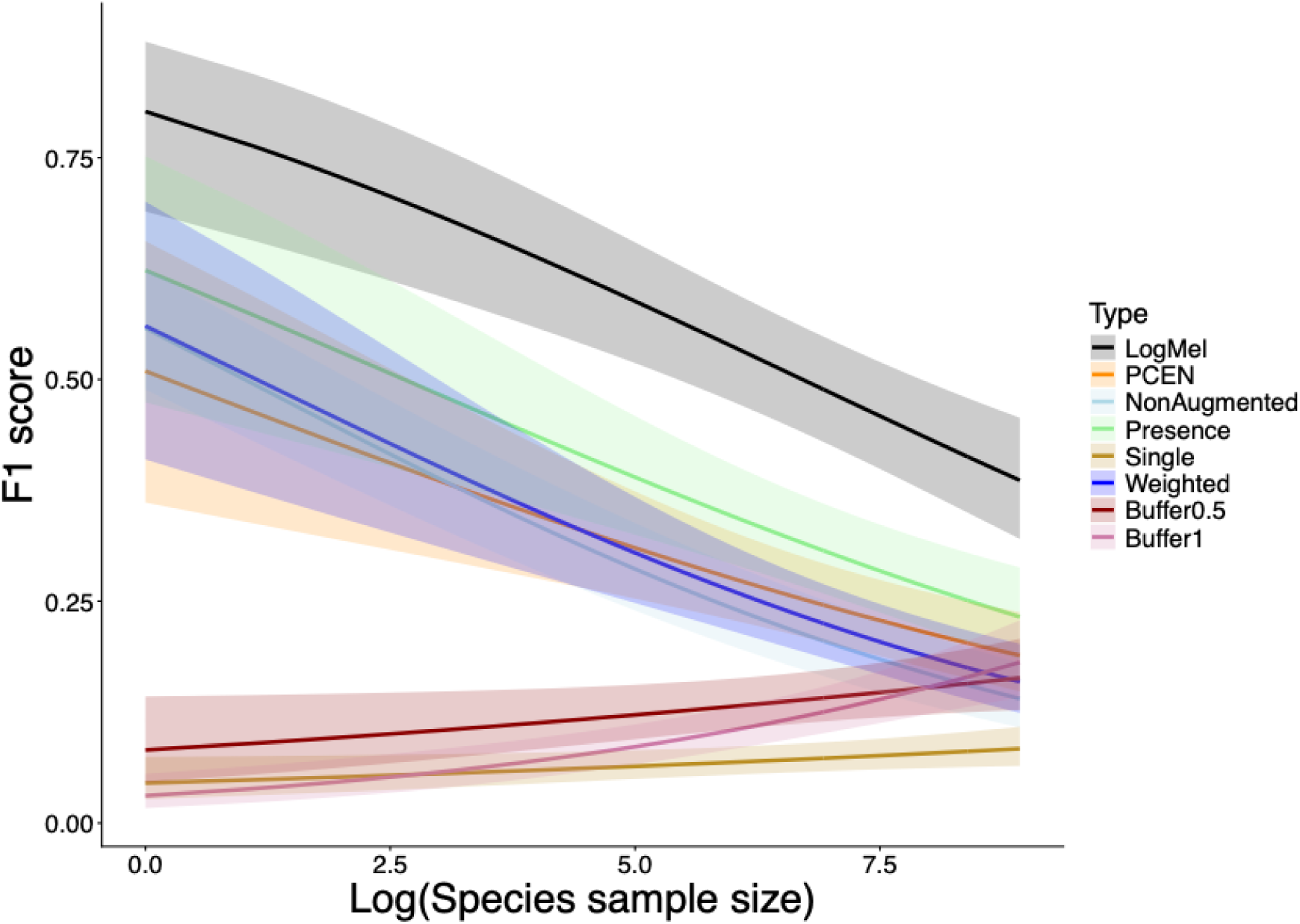
Interaction between model type and log-transformed sample size on predicted model performance assessed as F1-score (N = 10,584). Each line represents model-specific predictions from the fitted Generalized Additive Mixed Model, illustrating the interaction between model type and log-transformed sample size. Shaded areas indicate 95% confidence intervals.

Lastly, we found that model architecture significantly influenced CNN performance (Table 4). We specifically found that ConvNeXt outperformed the other evaluated CNN architectures in predicting species in tropical urbanization gradients (Table 4, Fig. 7). Underscoring the suitability of pretrained modern convolutional designs with enhanced feature representation capacity to accurately detect species in acoustically complex urban soundscapes when compared to older and not previously trained CNN architectures.

**Fig. 7.**
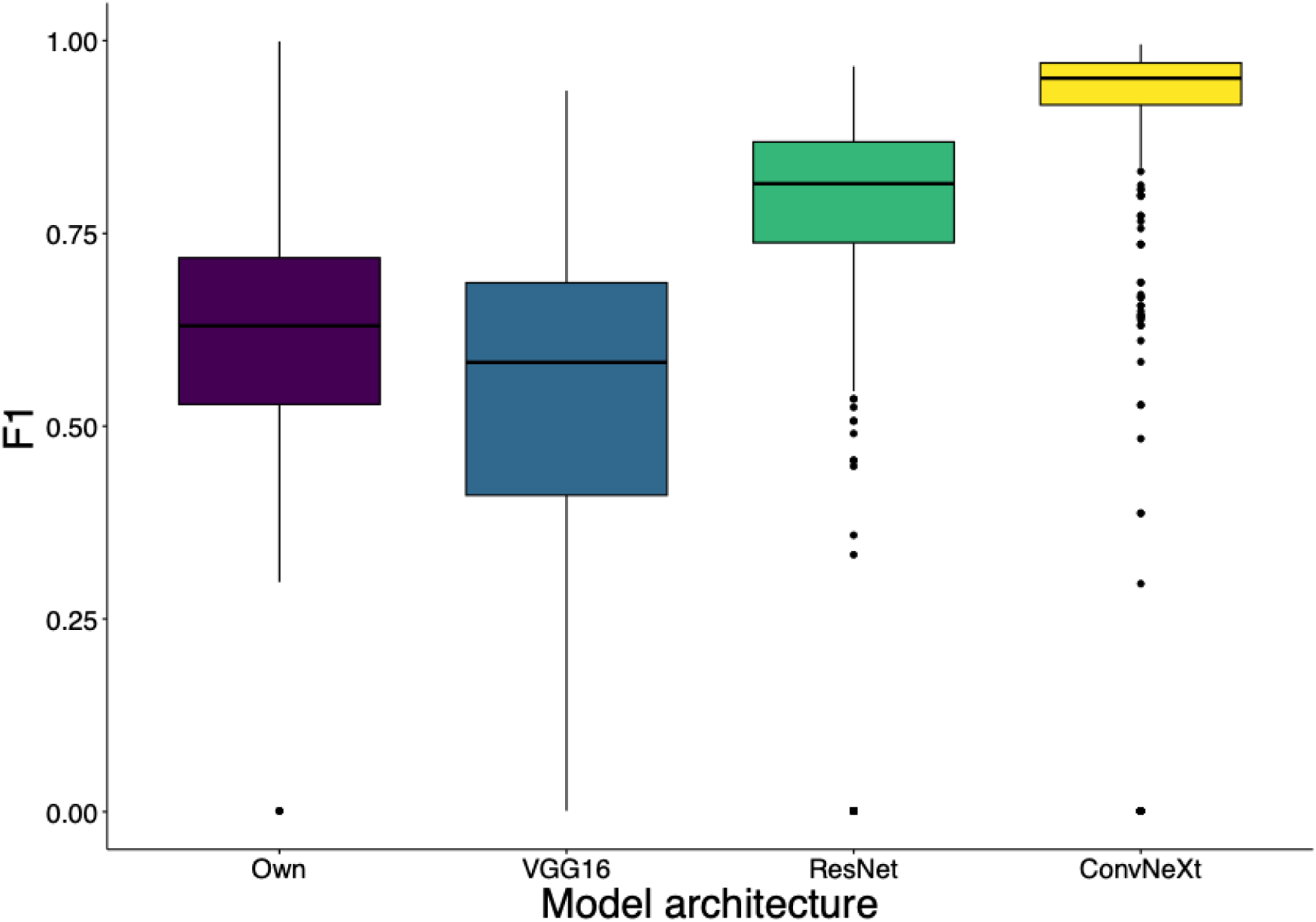
Differences of model performance, assessed as F1-score, across the different evaluated architectures. Each evaluated architecture is represented by a bar: Own (represented in dark purple, N= 441), VGG16 (in blue, N= 441), ResNet (in green, N= 441) and ConvNeXt (in yellow, N=441). The plot shows the medians (black line inside the box) with boxes showing the lower and upper quartile and black vertical bars showing the full range of the data. N= 1323 corresponds to the total data included in the model.

**Table 4.**
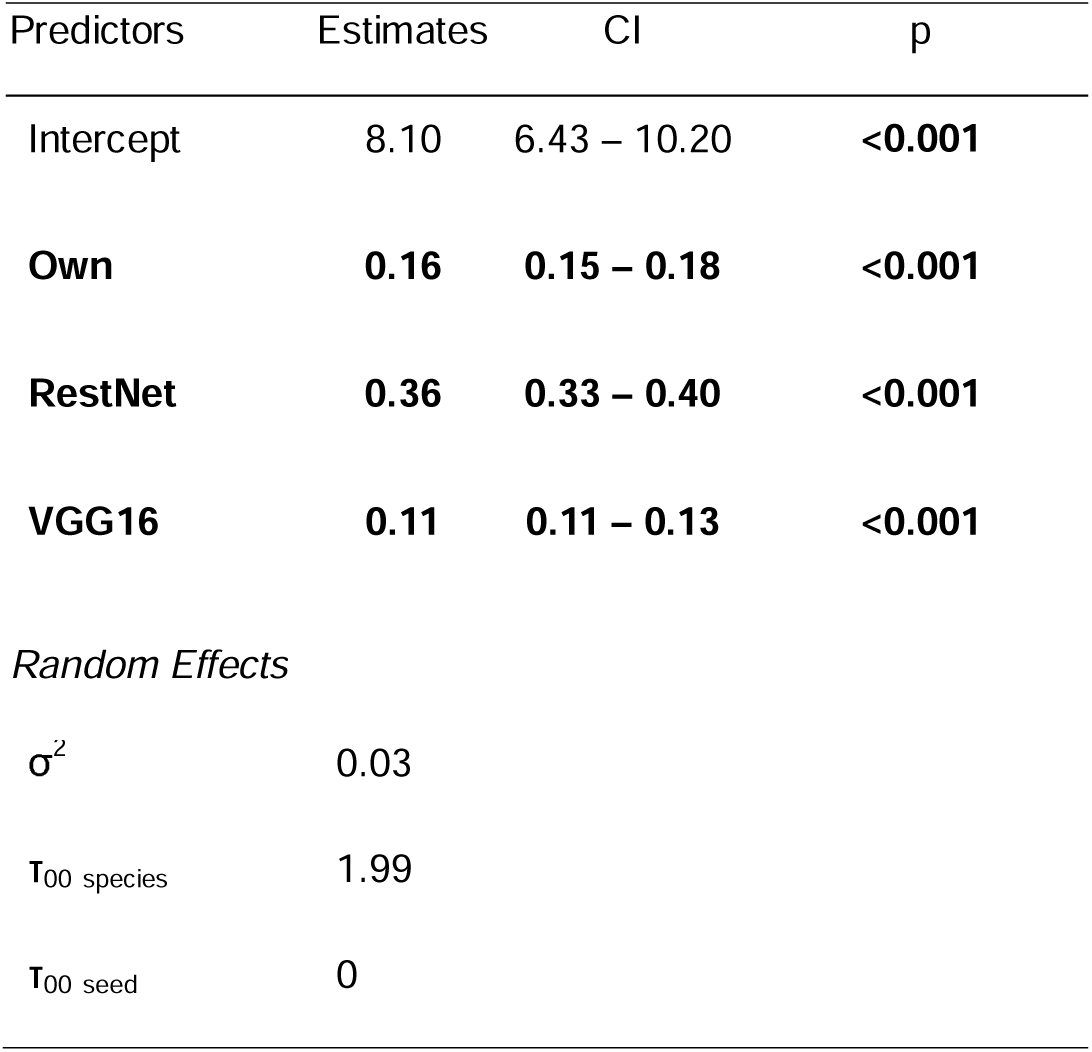
Coefficients, confidence intervals and p-values from a Generalized Linear Mixed Model built under the TMB template (glmmTMB) evaluating the influence of model architecture (fixed effect) on CNN performance (assessed as F1-score). The reference level for this model was the ConvNeXt architecture. Bold values indicate statistical significance at α = 0.05. *N* = 1323 corresponds to the total number of observations included in the regression model with 7 degrees of freedom. The variance explained by the fixed effect was *R*^2^ = 0.96 whilst the variance explained by both the fixed and random effects was *R*^2^ =0.04. Model AIC was -3493.408.

### 3.2 Comparing biodiversity estimates across monitoring methods

There was a high level of inter-observer agreement (72%) between the authors and the external expert in identifying the 42 species present in the evaluated recording, supporting the use of consensus-based species presence annotations as reference data. Using the reference annotations, we found that our CNN achieved higher micro and macro-averaged F1-scores than BirdNET for accurate species identification (Supplementary Table S5).

However, note that as this comparison was based on a single evaluation, no statistical testing was performed.

We found that species richness significantly differed between the CNN and point-count surveys (Table 5). Specifically, the CNN model detected a greater number of species than the point-count surveys (Table 5, Fig. 8). However, the two methods identified partially distinct species assemblages. Of the species detected by the CNN, 55 (53.922%) were recorded exclusively by this method and were not detected in the point-count surveys. In contrast, 43 species (51.19%) were recorded only during point-count surveys (Supplementary Table S6). These results indicate that, although the CNN approach achieved higher overall species detection in challenging ecosystems such as tropical cities, each method captured different components of the community.

**Fig. 8.**
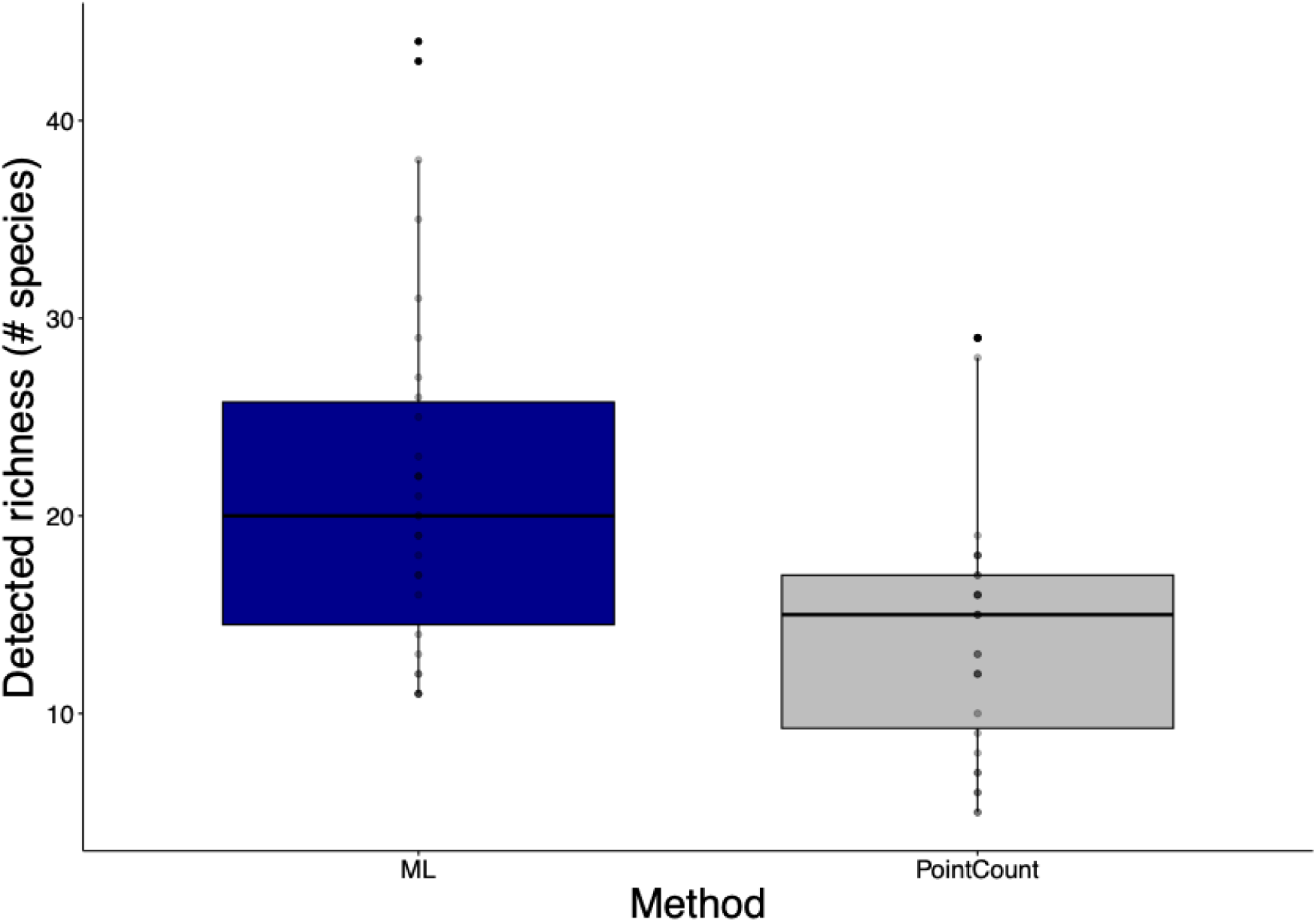
Differences between the richness detected between point-counts and Machine Learning (ML) approach. Each method is represented by a bar: ML (represented in dark purple, N= 30) and PointCount (represented in gray, N=30). The plot shows the medians (black line inside the box) with boxes showing the lower and upper quartile and black vertical bars showing the full range of the data. N= 60 corresponds to the total data included in the model.

**Table 5.** Estimate, standard errors and p-values from a Generalized Linear Model (GLM) evaluating the influence of method (fixed effect) on detected richness and the interaction between method and impervious surface. The reference level for this model was the CNN approach. *N* = 60 corresponds to the total number of observations included in the regression model. The variance explained by the fixed effect was *R*^2^ = 0.55. Bold values indicated statistical significance at α=0.05. Model AIC was 477.95.

| Predictors | Estimate | SD | z-value | p-value |
| --- | --- | --- | --- | --- |
| Intercept | 3.06 | 0.08 | 38.26 | $< 2 \times 10^{-16}$ |
| **Method (Point-Count)** | **-0.47** | **0.13** | **-3.68** | **0.000231** |
| ImperviousSurface | 0.02 | 0.13 | 0.14 | 0.89 |
| Method*ImperviousSurface | 0.1 | 0.21 | 0.49 | 0.63 |

Moreover, the interaction between method and impervious surface was not significant (Table 5), suggesting that the relationship between urbanization and richness was similar for the CNN and point-count methods, and therefore that urbanization affected species detection in comparable ways across both approaches.

Furthermore, we found a significant positive relationship between species richness derived from the best-performing CNN and the acoustic index NDSI, and a negative relationship with ADI (Table 6). Species richness increased as NDSI values increased, indicating that sites with greater biological acoustic activity relative to anthropogenic noise tended to exhibit higher CNN-derived richness (Fig. 9). In contrast, the negative relationship with ADI suggests that higher acoustic diversity values were associated with lower species richness, potentially reflecting the influence of broadband anthropogenic noise or other non-biological sounds that can inflate ADI in highly urbanized environments (Fig. 9). These results indicate that while NDSI may reliably reflect biodiversity patterns in urban tropical soundscapes, the relationship between ADI and biodiversity may vary depending on spectral diversity.

**Fig. 9.**
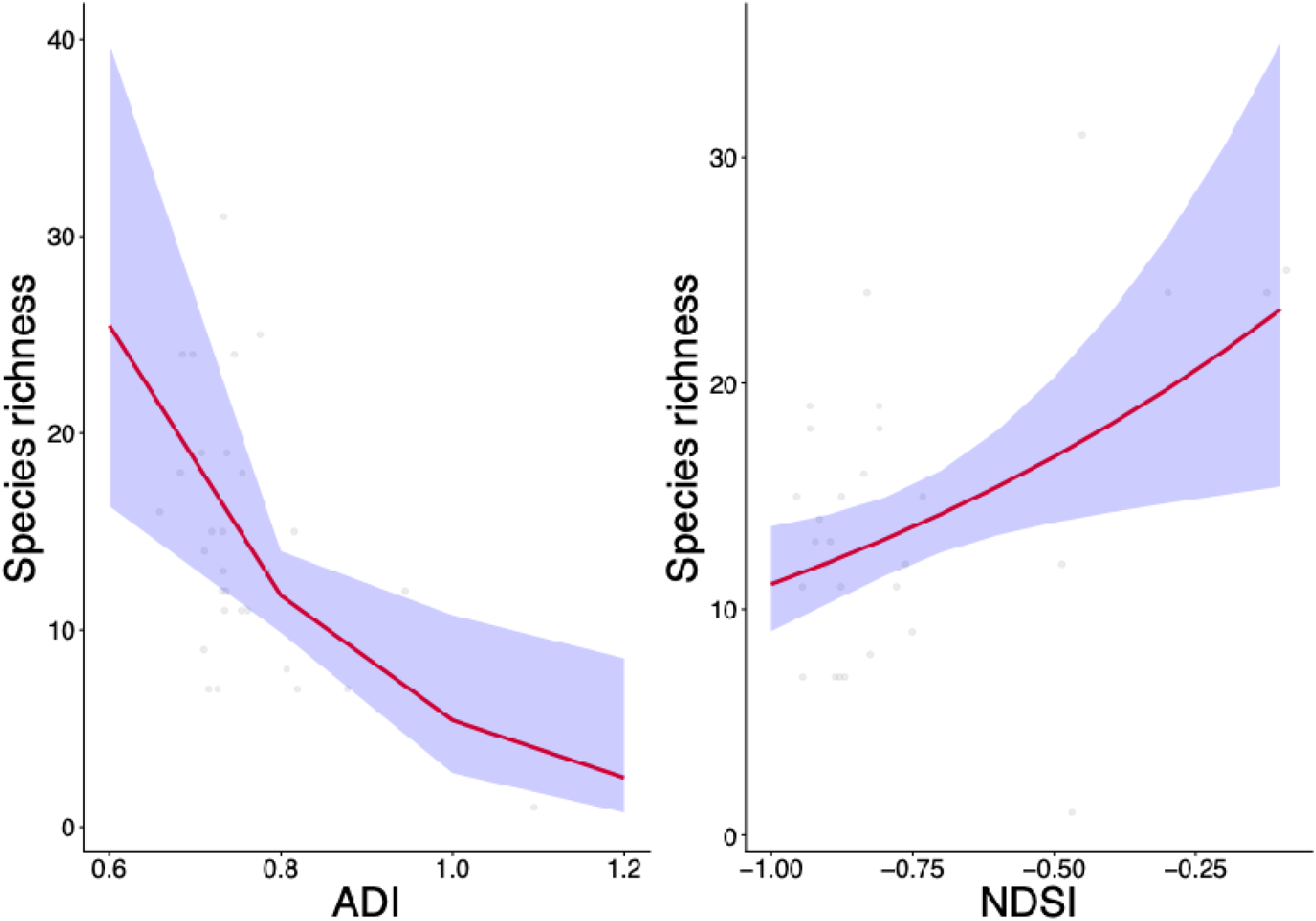
Relationship between the acoustic indices ADI (in the left panel) and NDSI (in the right panel) and species richness detected with our designed CNN (N = 30). The red line represents the predicted effect from the Negative Binomial Generalized Linear Model used to assess this relationship, while the blue shaded area indicates the corresponding 95% confidence interval.

**Table 6.** Coefficients, confidence intervals and p-values from a Negative Binomial Generalized Linear Model (GLM) evaluating whether acoustic indices captured biodiversity patterns similar to those detected by the CNN. Bold values indicate statistical significance at α = 0.05. *N* = 30 corresponds to the total number of observations included in the regression model with 1 degree of freedom. The variance explained by the fixed effect was *R*^2^ = 0.71. Model AIC was 185.66.

| Predictors | Estimate | SD | p |
| --- | --- | --- | --- |
| Intercept | 2.62 | 0.06 | **<0.001** |
| ACI | -0.02 | 0.07 | 0.738 |
| H | 0.01 | 0.12 | 0.942 |
| **ADI** | **-0.34** | **0.12** | **0.006** |
| **NDSI** | **0.2** | **0.08** | **0.009** |
| BI | -0.03 | 0.09 | 0.711 |

## 4. Discussion

As relatively few studies have simultaneously examined how model design choices influence CNN performance and how its biodiversity estimates compare with traditional monitoring approaches, our results highlight the importance of evaluating preprocessing and labelling strategies, probability thresholds, sample sizes, and model architectures when developing deep-learning tools, while also demonstrating that bioacoustic monitoring combined with CNN models can rapidly generate biodiversity estimates that detect a greater number of species than with traditional survey methods. Among the evaluated designs, CNN 2 (LogMel model) incorporating the ConvNeXt architecture, achieved the highest performance under a probability threshold of 0.3, suggesting that combining Log-Mel representations, multispecies labelling combining soundscape-specific training and external libraries data, modern pretrained architectures and larger training datasets can improve species identification in acoustically complex urbanization gradients in the Neotropics. Furthermore, comparisons of biodiversity estimates across monitoring approaches showed that automated acoustic classification achieved higher species detections than traditional point counts, although the two methods captured partially distinct species assemblages. In addition, positive relationships between CNN-derived richness and acoustic indices such as NDSI suggest that this index can potentially capture similar biodiversity patterns and may serve as useful and rapid proxies for biodiversity in acoustic monitoring. Together, our findings demonstrate that 1) model design, threshold selection, architecture, and data availability jointly shape CNN performance and should be carefully considered during model development, and 2) automated acoustic classification can provide rapid biodiversity assessments that yield higher species detection rates compared to traditional monitoring approaches, highlighting its potential to support biodiversity monitoring in complex tropical urbanized ecosystems.

### 4.1 Identification of the best CNN model

#### 4.1.1 Identification of the best model design

The observed performance improvements associated with the use of Log-Mel spectral representations, the inclusion of presence and absence labels, and multispecies field and external libraries recordings without buffer zones or species weighting suggests that effective CNN performance in acoustically complex ecosystems depends on: 1) applying spectral transformations that highlight biologically relevant acoustic features, 2) using training data that combine recordings resembling real soundscape conditions with cleaner reference vocalizations, 3) avoiding non-informative labelling strategies that can hinder learning, and 4) recognizing that species weighting may not address the primary challenges posed by acoustically complex datasets. Overall, these findings highlight that the choice of spectral representation, labelling strategies and choice of including species weighting should be guided by the acoustic characteristics of the soundscapes and targeted species.

First, contrary to our initial prediction, we found that Log-Mel yielded the best CNN performance. We suggest that Log-Mel spectral representations outperformed PCEN because they better preserve frequency components critical for species identification, particularly the low to mid-frequency ranges where most bird vocalizations occur, that can be partially attenuated by energy-normalizing transformations such as PCEN (Dias et al., 2025). Our findings are consistent with previous studies showing that Log-Mel spectrograms yielded stronger performance for bird species identification in Brazil (Dias et al., 2025) and for buzz-pollinating bee species in Chile (Ferreira et al., 2025). Although other study reported superior performance of PCEN for general sound classification tasks (Lostanlen et al., 2019a), this advantage may not directly translate to species-level classification, where fine spectral details are critical for distinguishing acoustically similar species, as demonstrated by our results. Finally, due to Mel-scale representations being originally designed to simulate human auditory perception (Logan, 2000), their suitability for species identification may vary across taxa, highlighting that the choice of spectral representation should be guided by the acoustic characteristics of the targeted taxa (as also suggested by Dias et al. 2025).

Second, the improved performance of CNNs trained with both presence and absence data, together with multispecies labels, supports the idea that training labels closely reflecting real soundscape conditions enhance model learning in acoustically complex ecosystems (Knight et al., 2017). Providing multispecies annotations and explicit absence data enables models to better distinguish target species vocalizations from other soundscape elements, as well as to identify species within overlapping vocalizations reducing false positives and improving discrimination in urban soundscapes, which is consistent with previous studies showing that incorporating acoustically realistic training data enhances model performance (Knight et al., 2017; Lostanlen et al., 2019a; Priyadarshani et al., 2018). Together, these findings highlight the importance of aligning labelling strategies with the ecological and acoustic complexity of the target ecosystem.

The importance of using training data combining vocalizations closely resembling the target ecosystem and cleaner reference recordings that capture geographic variation was supported by the improved performance of models trained on both field and XenoCanto recordings. Recordings from online sound libraries such as XenoCanto, are often single-species vocalizations with limited background noise, providing clearer acoustic examples from which models can learn from (as suggested by Clark et al., 2023). Additionally, the XenoCanto recordings used in this study were sourced from different countries across the Neotropical region, potentially increasing the acoustic variability represented within species and thus, improving model flexibility to geographic variation in vocalizations (Clark et al., 2023). Contrastingly, field recordings can reflect better the complexity of real soundscapes, including overlapping vocalizations and environmental noise. Together, these complementary data sources may have improved model performance by exposing CNNs to both acoustically clean reference signals that capture geographic acoustic variation and recordings that closely reflect soundscape conditions. This pattern is consistent with previous studies showing that training CNNs with external vocalizations form XenoCanto improved model performance in both acoustically complex environments (Clark et al., 2023) and less-disturbed ecosystems (Schiavo et al., 2025), suggesting that integrating cleaner training data may improve CNN robustness across heterogeneous soundscapes. Nonetheless, previous work has also shown that incorporating recordings from online sound libraries can create domain mismatches between training data and real-world soundscapes with CNN classification accuracy often improving when training data originates directly from the study system (Lauha et al., 2022; MacIsaac et al., 2024). Discrepancies between those studies and our findings may be related to differences in training dataset size, as our field-recording dataset was comparatively limited for some species (approximately 10-5075 recordings per species; Supplementary Table S3), Consequently, incorporating additional samples may have provided greater acoustic diversity and clearer representations, thereby improving CNN performance. These results suggest that including additional samples from external libraries might be beneficial when data tends to be limited but might become redundant once dataset size and acoustic variability are sufficiently represented. Lastly, data augmentation can also be achieved with strategies that transform the original recordings such as time stretching, pitch shifting, noise injection, and spectrogram augmentation that were not explored in this study but have been shown to improve CNN performance in acoustic classification tasks, could potentially improve its accuracy in acoustically complex ecosystems (as suggested by Nanni et al. 2020). Therefore, future research should assess whether such augmentation strategies can enhance CNN performance for bird species identification in urban tropical ecosystems.

Padding labels (i.e., adding buffer zones) not improving the performance of CNNs is likely due to the inclusion of non-informative diluting species-specific features by introducing label noise, thereby reducing the ability of the models to learn discriminative patterns for each species (as suggested by Tang et al. 2019). Label noise affecting model performance is coherent with previous studies finding that the inclusion of non-informative sections (such as silence buffer zones) hinders model performance (Ding et al., 2022; Karimi et al., 2020). Nonetheless, other models such as the widely used BirdNET have found that adding context frames around calls (i.e., buffer zones) improve their performance (Kahl et al., 2021), suggesting that the effectiveness of zero padding in CNNs is likely ecosystem-dependent, with padding providing useful contextual information in some acoustic environments but introducing label noise in others, such as highly complex urban tropical soundscapes.

Contrary to our initial expectations, even though our dataset was highly unbalanced across species (Supplementary Table 3), species weighting did not improve CNN performance, suggesting that class imbalance might not have been the primary limiting factor in our dataset. In acoustically complex urban soundscapes, other factors such as overlapping vocalizations, background noise, and interspecific acoustic similarity can be more constraining for identification than sample size alone (as suggested by Priyadarshani et al. 2018). Indeed, in such contexts, class weighting can push the model to emphasize rare species, but when species share similar acoustic features or vocalize together with background noise, this may not help the model learn clearer distinctions and thus, reduce overall performance (Priyadarshani et al., 2018). Moreover, the use of multispecies labels in our models may have already facilitated learning for rare species, thereby reducing the need for additional species weighting. Lastly, although class weighting is commonly applied in acoustic identification models to address class imbalance and has been shown to improve performance in some contexts (Elfwing et al., 2018; Razali et al., 2025; Sothe et al., 2020), similar findings in ecological and machine learning studies have shown that weighting strategies do not often offer benefits when classes overlap and errors are driven mainly by acoustic similarity, overlapping vocalizations, and background noise rather than class imbalance (Priyadarshani et al., 2018; Stowell et al., 2019a; Xiong et al., 2026). Thus, these results suggest that the inclusion species weighting should be justified based on the ecological context and the structure of the data, as its effectiveness can be context dependent.

We found that CNNs performed best when using a probability threshold of 0.3. However, using this relatively low threshold involves a trade-off between precision and recall, where improving the model sensitivity is achieved at the expense of increasing false positives (Miller et al., 2024). Nonetheless, in acoustically complex urban tropical environments where bird vocalizations are often weak and partially masked by background noise or overlapping species, a lower detection threshold can help recover true positives that would otherwise be missed under more conservative settings (Priyadarshani et al., 2018). Although some studies have reported improved performance at higher thresholds (e.g., 0.5), these findings are typically associated with less acoustically complex ecosystems (Pérez-Granados et al., 2025). In contrast, a threshold of 0.3 is commonly adopted in models designed to operate under challenging soundscape conditions with simultaneous vocalizations, such as BirdNET (Kahl et al., 2021), supporting its suitability for urban environments. Together, these findings suggest that optimal threshold selection is strongly influenced by both acoustic complexity and species richness, with lower thresholds being more suitable for bird detection in noisy, species-rich environments such as urban tropical ecosystems, whereas higher thresholds may be more appropriate in less noisy and diverse acoustic settings (adapted from Miller et al. 2024; Pérez-Granados et al. 2025). However, further studies assessing how CNN performance varies across different probability thresholds are needed to better understand how threshold selection influences the balance between precision and recall across ecosystems and monitoring objectives.

Furthermore, although model performance generally improved as more training data became available, these improvements tended to plateau at higher sample sizes. Model performance plateauing at higher samples might reflect feature saturation, a common pattern in machine-learning, where beyond a certain threshold additional recordings contribute limited new acoustic information and may therefore become increasingly redundant for model training (Hestness et al., 2017). This finding is consistent with previous work showing that CNN classification performance can eventually stabilize once datasets become sufficiently large (Hestness et al., 2017; Lauha et al., 2022). However, previous studies have contrastingly showed that increasing training data can improve CNN classification accuracy in a model designed to identify 18 small-mammal species (MacIsaac et al., 2024) and a presence-only species distribution CNN (Anand et al., 2026). Suggesting that dataset size may be an important predictor in the relationship between sample size and model performance, as increasing sample size can substantially improve CNN performance when data are limited, but cause data saturation once datasets become sufficiently large (Hestness et al., 2017). Additionally, the magnitude of performance improvements varied across model designs, potentially reflecting differences in learning dynamics and how efficiently each architecture benefits from additional training data (Rosenfeld et al., 2019). Some models may reach performance saturation more rapidly, whereas others continue to improve with increasing sample size (Rosenfeld et al., 2019). These findings suggest that, although sample size can be an important factor influencing CNN performance, it is not sufficient on its own; data quality, acoustic representativeness and the optimization of model architecture and training strategies may be as equally important.

Lastly, our results showed that the random-effects structure, accounting for variation among species and training seeds, explained a larger proportion of variance than the model designs alone. Indicating that substantial heterogeneity in model performance is driven by species-specific characteristics and stochastic variation associated with model initialization and training rather than solely by the different preprocessing and labelling strategies evaluated. Species-level differences likely reflect variation in vocalization structure, detectability and acoustic similarity between species and other soundscape elements that can strongly influence CNN performance (Priyadarshani et al., 2018; Stowell et al., 2019a). These effects may be particularly relevant in our study which encompasses urbanization gradients where soundscape composition varies considerably due to differences in the number and identity of vocalizing species, the type and intensity of anthropogenic noise and their combined influence on the acoustic environment. Similarly, variation among seeds highlights the sensitivity of deep learning models to stochastic optimization processes, particularly in complex and imbalanced datasets (Bouthillier et al., 2021). Together, these findings suggest that species identity can be a key determinant of model performance and that training stochasticity can meaningfully influence results, underscoring the importance of accounting for both species-level variability, representation of different types and intensities of anthropogenic noise and training randomness when evaluating and comparing CNN performance (Bouthillier et al., 2021; Stowell et al., 2019a). Consequently, future studies should report results across multiple training runs and explicitly evaluate species and soundscape-specific performance to ensure robust and reproducible CNN models.

#### 4.1.2 Identifying the best model architecture

We found that the highest CNN performance was achieved by a pretrained ConvNeXt model incorporating the optimal modelling design. ConvNeXt has been shown to offer enhanced hierarchical feature representation and more efficient optimization by incorporating modern design elements such as larger convolutional kernels and simplified architectural blocks (Liu et al., 2022), which may allow it to better capture the subtle, overlapping acoustic patterns characteristic of urban tropical soundscapes, explaining its superior performance relative to models trained from scratch and those based on older CNN architectures. Nonetheless, ConvNeXt is a relatively recent architecture, and studies comparing its performance with established backbones such as VGG16 and ResNet are limited. However, generally, most studies have shown that ResNet architectures frequently outperform shallower networks in bird sound classification such as VGG16, as residual connections enable the learning of deeper and more robust feature hierarchies from spectrograms (Azhari et al., 2025; LeBien et al., 2020). However, VGG-16 has also achieved high accuracy in bird species identification when combined with Log-Mel spectrograms and transfer learning (Madake et al., 2023), highlighting that different architectures can perform well depending on dataset characteristics and preprocessing strategies (as suggested by Dey et al. 2026). Specifically, VGG-16 may be better suited to less acoustically complex ecosystems dominated by single-species vocalizations (Madake et al., 2023), whereas ResNet architectures often perform well in moderately noisy environments due to their ability to learn deeper feature hierarchies (LeBien et al., 2020) and, ConvNeXt appears especially effective in highly complex soundscapes, such as urbanized tropical environments, where overlapping vocalizations and background noise require optimized representations. Together, these findings suggest that architectural performance can be strongly context dependent and closely linked to soundscape characteristics and that, while ConvNeXt represents a promising advancement for bird acoustic identification in complex soundscapes, further studies are needed to systematically assess its efficiency and generalizability across a wider range of ecosystems and acoustic conditions.

### 4.2 Comparing biodiversity estimates across monitoring methods

#### 4.2.1 Comparison between CNN and BirdNET performance

When comparing the inferences generated by our CNN with those produced by BirdNET using ground-truth annotations independently provided by an expert and by us, we found that our CNN achieved higher performance, however, note this comparison was based on a single evaluation and was therefore not statistically tested thus, the explanations provided below should be considered tentative. Our CNN yielding a better performance could be a result of BirdNET been primarily trained on data from temperate regions, which can limit its ability to identify species in distinct tropical ecosystems such as ours. BirdNET providing a limited performance in tropical ecosystems is consistent with a global assessment of BirdNET performance across biomes, reporting lower predictive performance in Central and South America (PR AUC ≈ 0.13) and Africa and Asia (PR AUC ≈ 0.03–0.04), compared with temperate regions (PR AUC ≈ 0.16–0.23) (Funosas et al., 2026). However, other study reported high BirdNET performance for identifying a three tropical species (Pérez-Granados and Schuchmann, 2025); nonetheless, this result may be attributable to the lower number of target species assessed and the monitoring being performed in a natural ecosystem with less acoustic challenges for the identification of species. Together, these findings suggest that developing and training ecosystem-specific models may be advantageous when working with acoustic data from tropical regions, where high species richness, frequent acoustic overlap, and distinct vocalization structures may not be fully captured by globally trained models however, future studies incorporating multiple comparisons will be needed to robustly assess differences between the performance of BirdNET and ecosystem-specific CNN models.

#### 4.2.2 Comparing richness across our CNN, point-counts and acoustic indices

Following our initial predictions and the commonly reported limitations of point-count surveys, CNN-derived approaches were able to detect more species than traditional point counts. The higher species detection achieved by CNN models may be explained by the broader temporal coverage provided by autonomous acoustic recorders (Darras et al., 2019), which sampled continuously over a week, whereas point counts consisted of only two 10-minute surveys per site. Extended acoustic monitoring increases the likelihood of detecting species with varying vocal activity patterns (Darras et al., 2019). In addition, automated acoustic approaches can reduce observer-related biases and allow repeated examination of recordings, potentially improving the detection of species overlooked during field observations (Darras et al., 2019, 2018). Similar patterns have been reported in previous studies comparing autonomous acoustic monitoring and traditional surveys that reported higher species detection with automated recorders (Doohan et al., 2026; Schuster et al., 2024). However, evidence regarding which method is more effective in species detection remains mixed. Some studies report similar richness between visual and acoustic approaches in Mount Kenya (wa Maina and Njoroge, 2025), and Brazilian open woodlands (Alquezar and Machado, 2015), likely reflecting the complementary strengths of both methods as whilst point counts integrate visual and auditory cues, acoustic monitoring relies solely on vocal detections but enables continuous sampling over longer periods (Darras et al., 2018). In other cases, studies find higher richness with either point counts (Leach et al., 2016; Stevens et al., 2019) or acoustic recorders (Doohan et al., 2026; Schuster et al., 2024), suggesting that outcomes depend on species detectability and community composition (as suggested by Alldredge et al. 2007; Farmer et al. 2012). Lastly, impervious surface affected both methods similarly, suggesting that although CNN-based acoustic monitoring detected a greater number of species, both approaches captured comparable ecological responses to urbanization gradients as also reported in previous studies showing that automated recorders and point-counts often detect consistent ecological responses despite differences in overall species richness and detectability (Budka et al., 2021; Darras et al., 2018).

Additionally, differences in detectability between methods may not only influence their relative efficiency but also the composition of the communities detected (as suggested by Darras et al., 2018). In our study, both methods detected partially distinct assemblages with point-count surveys more frequently detected less vocal species such as doves and generally, species that tend to be more tolerant to human presence, whereas acoustic approaches more commonly detected species that may be less tolerant to humans such as antpittas or more easily identified through vocal activity due to their physical similarity such as certain flycatchers (Supplementary Table S6). This pattern is consistent with previous studies from Mount Kenya (Wa Maina and Njoroge, 2025) and Brazil (Alquezar and Machado, 2015), where acoustic and visual survey approaches detected partially distinct species assemblages potentially due to each method capturing different components of bird communities. Together, our results highlight the potential of acoustic approaches combined with deep learning methods such as CNNs to monitor avian diversity in acoustically challenging ecosystems, while also suggesting that integrating visual surveys with automated acoustic monitoring may provide a more comprehensive assessment of bird community composition and biodiversity patterns.

Moreover, although the CNN was trained on 147 species identified during the labelling phase, the model only detected 103 species across the study sites. Because species labels were derived from our dataset, this discrepancy likely reflects differences in detectability between the manually labelled subset and the full acoustic dataset, as active listening during the labelling phase allowed us to detect species that may have been missed by the CNN due to infrequent vocalizations, low-amplitude calls or masking by environmental and anthropogenic noise. Such differences in detectability between labels and acoustic datasets highlight a common limitation of automated acoustic approaches, where rare, quiet or acoustically complex vocalizations may be under-detected compared to manual inspection (Gibb et al., 2019). Similar patterns have been reported in previous studies applying multi-species acoustic classifiers, where models trained on large labelled datasets only detected a subset of those species when applied to specific soundscapes (Kahl et al. 2021; Stowell et al. 2019), suggesting that local soundscape conditions and species vocal behavior could shape acoustic detection however, more studies are needed to establish this. Some approaches such as integrating automated detection with manual verification and expanding training datasets with rare and low-amplitude vocalizations could help improve detection of quiet or acoustically complex calls (Priyadarshani et al., 2018; Stowell et al., 2019a) and thus, should be considered in future studies to improve the performance of CNN in these soundscapes.

Although acoustic indices are widely used as proxies for biodiversity, their performance in urban ecosystems remains inconsistent (Fairbrass et al., 2017; Santos et al., 2024). Among the evaluated indices, we found that the Acoustic Diversity Index (ADI) showed a negative relationship with species richness inferred by our best-performing CNN, whereas the Normalized Difference Soundscape Index (NDSI) showed an inverse relationship. First, ADI measures the distribution of acoustic energy across frequency bands (Villanueva-Rivera et al., 2011), which is often expected to increase as more species vocalize within different acoustic niches. However, ADI reflects the acoustic diversity of all sound sources including biological, geological, and anthropogenic signals. As a result, it can capture spectral heterogeneity generated by non-biological soundscape elements, which makes its interpretation particularly challenging in urban ecosystems where these sources often occur simultaneously (revised in Xu et al. 2023). Consequently, sites with high levels of anthropogenic noise or other urban disturbances may exhibit inflated ADI values that do not necessarily reflect true biological diversity (Xu et al., 2023), resulting in higher ADI levels in sites with low diversity. Alternatively, urbanization can also lead to the homogenization of avian communities where to avoid signal masking by urban noise, bird species may adjust their vocalizations to similar frequency ranges (Hao et al., 2024). As a result, even when multiple species are present at a site, their calls may overlap within similar frequency bands (Slabbekoorn and Peet, 2003), potentially producing lower ADI values despite relatively high species diversity (suggested in Alcocer et al. 2022). Therefore, although ADI may not be fully reliable for biodiversity monitoring, it could still provide insights into how urbanization influences bird vocalization frequencies however, future research should examine how soundscape components such as anthropogenic noise, community composition, and urban-driven shifts in vocalization frequency affect ADI to better understand its suitability in urban contexts (suggested by Alcocer et al. 2022).

Secondly, NDSI reflects the balance between biological and anthropogenic sounds, with higher values occurring when biological acoustic activity dominates over anthropogenic noise (Kasten et al., 2012). Consequently, sites with greater vocal activity and lower anthropogenic disturbance tend to exhibit higher NDSI values and greater species richness (Kasten et al., 2012), which may explain their correspondence with CNN-derived biodiversity estimates. Additionally, because NDSI directly incorporates the relative contribution of biological and anthropogenic acoustic energy (Kasten et al., 2012), increases in species richness and associated vocal activity can elevate the biological component of the soundscape, further strengthening the relationship between this index and biodiversity. The positive association between NDSI and biodiversity is consistent with previous studies in an urban nature reserve in Brazil (Alquezar and Machado, 2015) and urban parks in Iran (Latifi et al., 2023), where this acoustic index associated with bird species richness, suggesting its potential as a biodiversity proxy in challenging soundscapes. Contrastingly, as mentioned before, information regarding whether acoustic indices could be applied under urban contexts remains mixed with some studies have reported a lack of association between species richness and acoustic indices in urban parks in Brazil (Santos et al., 2024) and London (Fairbrass et al., 2017), suggesting that the relationship between acoustic indices and biodiversity can vary depending on local environmental conditions, species composition, and background noise levels and thus, future studies should consider all these factors before applying these indices in there study system.

Moreover, given the ability of each index to capture distinct aspects of soundscapes, combining different indices could increase their potential for monitoring biodiversity as reported in previous studies (Buxton et al., 2018; Towsey et al., 2014). However, further research is needed to determine the effect of combining acoustic indices as biodiversity proxies in urbanized ecosystems. Lastly, although acoustic indices can provide rapid biodiversity assessments, they are limited in their ability to capture species-level information such as community composition (Giuliani et al., 2024); therefore, studies aiming for more detailed biodiversity assessments should consider using them as complementary tools alongside species-level monitoring approaches such as automated acoustic classification.

## 5. Conclusion

In conclusion, this study showed that CNN performance for bird identification in tropical urban gradients depends strongly on model design and architecture. CNN performance improved with Log-Mel spectrograms, training data combining field and XenoCanto recordings, appropriate probability thresholds and a ConvNeXt backbone, whereas label padding and species weighting did not improve performance in this context. Although larger sample sizes generally increased performance, it tended to plateau at higher sample size and patterns varied across models. Moreover, our locally trained CNN outperformed BirdNET, suggesting that region-specific models may be better suited to acoustically distinct tropical ecosystems. Beyond model optimization, CNN models were able to detect more species than traditional point-counts, highlighting the potential of automated acoustic monitoring for biodiversity assessment. In addition, NDSI was associated with CNN-derived richness, supporting its use as a rapid biodiversity proxy, whereas ADI may still provide complementary information on how urbanization affects bird vocal activity. Overall, our findings emphasize the importance of tailoring CNN design to the target ecosystem and show that automated acoustic classification can enhance biodiversity monitoring in complex tropical urban ecosystems by detecting a broader range of species while complementing traditional surveys and acoustic indices. Future work should combine automated detection with manual validation and expand training datasets to improve the detection of rare or low-amplitude vocalizations.

## Declaration of competing interest

The authors declare that they have no known competing financial interests or personal relationships that could have influenced this work.

## Acknowledgements

We want to thank Diba Andrade, Miguel Moreno-Palacios and Sebastian Betancourth for their generous support during fieldwork and, the universities, regional and local institutions, reserves, and communities that allowed us to conduct visual surveys and deploy acoustic recorders in their territories. We also thank Daniel Sánchez for helping during data collection and Jaime Andres Garizabal Carmona for generously annotating a dataset used to assess the performance of our CNN. We also thank the editor and reviewers for their thoughtful feedback, which strengthened the quality of this work.

## Data availability

Code used to train, validate and test the best performing model, can be accessed in DOI: 10.17632/5s35783d2t.1

Data will be made available on request.

## Notes

### Competing Interest Statement

The authors have declared no competing interest.

### Summary of Updates

Results and Discussion were updated from last version. Supplementary material is now available

