## Supplementary Material for "Tuning into the city soundscape: Optimizing Convolutional Neural Networks for avian acoustic identification in the neotropics and evaluating their performance against established monitoring approaches"

***Table S1.*** Summary of principles, examples and outcomes of the most commonly used acoustic indices to assess biodiversity in different ecosystems.

| Acoustic index | Ecological Principle | Study | Ecosystem | Result |
| --- | --- | --- | --- | --- |
| **BIO**: Bioacoustic index  (Boelman et al. 2007) | Quantification of the total area under the curve of sounds amplitude. This measurement indicates the extent of natural sounds (i.e., biophony) relative to human noise (anthropophony). | (Mitchell et al. 2025)  (Gaspar et al. 2023)  (Diaz et al. 2023a)  (Kotian et al. 2024)  (Alcocer et al. 2022)  (Fairbrass et al. 2017)  (Jorge et al. 2018)  (Santos et al. 2024) | Temperate natural  Tropical natural  Tropical urban  Tropical natural  Both natural  Temperate urban  Tropical natural  Tropical urban | Accurate  Accurate  Not accurate  Accurate  Not accurate  Accurate  Not accurate  Not accurate |
| **H:** Entropy Index  (SUEUR et al. 2008) | Measurement of relative sound energy abundance of the amplitude of sounds across frequency bands (i.e., evenness), reflecting the richness of the soundscape. | (Gaspar et al. 2023)  (Diaz et al. 2023a)  (Kotian et al. 2024)  (Alcocer et al. 2022)  (Devos 2023)  (Dröge et al. 2021)  (Jorge et al. 2018)  (Santos et al. 2024) | Tropical natural  Tropical urban  Tropical natural  Both natural  Temperate urban  Tropical natural  Tropical natural  Tropical urban | Not accurate  Not accurate  Not accurate  Accurate  Accurate  Accurate  Not accurate  Not accurate |
| **ACI:** Acoustic complexity index  (Pieretti et al. 2011) | Assessment of variability of sounds energy within band frequencies across short intervals of time. | (Mitchell et al. 2025)  (Kotian et al. 2024)  (Diaz et al. 2023a)  (Alcocer et al. 2022)  (Fairbrass et al. 2017)  (Devos 2023)  (Dröge et al. 2021)  (Jorge et al. 2018)  (Santos et al. 2024) | Temperate natural  Tropical natural  Tropical urban  Both natural  Temperate urban  Temperate urban  Tropical natural  Tropical natural  Tropical urban | Accurate  Accurate  Accurate  Accurate  Accurate  Accurate  Not accurate  Not accurate  Not accurate |
| **ADI:** Acoustic diversity index  (Villanueva-Rivera et al. 2011) | Assessment of the spread of sounds energy across energy within a recording, indicating its complexity. | (Mitchell et al. 2025)  (Gaspar et al. 2023)  (Diaz et al. 2023a)  (Alcocer et al. 2022)  (Fairbrass et al. 2017)  (Devos 2023)  (Dröge et al. 2021)  (Machado et al. 2017)  (Jorge et al. 2018) | Temperate natural  Tropical natural  Tropical urban  Both natural  Temperate urban  Temperate urban  Tropical natural  Tropical urban  Tropical natural | Accurate  Not accurate  Not accurate  Not accurate  Not accurate  Accurate  Accurate  Accurate  Not accurate |
| **AEI:** Acoustic evenness index  (Villanueva-Rivera et al. 2011) | Uses the Gini Index to quantify the evenness of sounds energy across different frequencies within the recording. | (Gaspar et al. 2023)  (Diaz et al. 2023a)  (Alcocer et al. 2022)  (Devos 2023)  (Dröge et al. 2021)  (Jorge et al. 2018) | Tropical natural  Tropical urban  Both natural  Temperate urban  Tropical natural  Tropical natural | Not accurate  Not accurate  Not accurate  Accurate  Accurate  Accurate |
| **AR:** Acoustic richness  (Depraetere et al. 2012) | Uses the diversity of amplitude and time entropy as a proxy for biodiversity. Thus, reflecting how consistent high-amplitude sounds are within a recording. | (Gaspar et al. 2023)  (Diaz et al. 2023a)  (Alcocer et al. 2022) | Tropical natural  Tropical urban  Both natural | Not accurate  Not accurate  Not accurate |
| **NDSI:** Normalised difference soundscape index  (Kasten et al. 2012) | Measure of anthropophony in a soundscape by comparing acoustic energy from biophony and anthropophony. | (Mitchell et al. 2025)  (Gaspar et al. 2023)  (Kotian et al. 2024)  (Alcocer et al. 2022)  (Fairbrass et al. 2017)  (Devos 2023)  (Machado et al. 2017)  (Jorge et al. 2018)  (Santos et al. 2024) | Temperate natural  Tropical natural  Tropical natural  Both natural  Temperate urban  Temperate urban  Tropical urban  Tropical natural  Tropical urban | Accurate  Not accurate  Not accurate  Accurate  Accurate  Accurate  Not accurate  Not accurate  Not accurate |
| **NP:** Number of frequency peaks  (Gasc et al. 2013) | Counts the number of peaks in a frequency spectrum over a time period to quantify acoustic activity. | (Kotian et al. 2024)  (Alcocer et al. 2022) | Tropical natural  Both natural | Not accurate  Not accurate |

***Fig. S1.*** Map highlighting the selected study sites in Bogotá, Medellín, Ibagué, Tunja and Neiva. Sites represented an urbanization gradient and are indicated by red points. This image was recovered from Esri World Imagery.

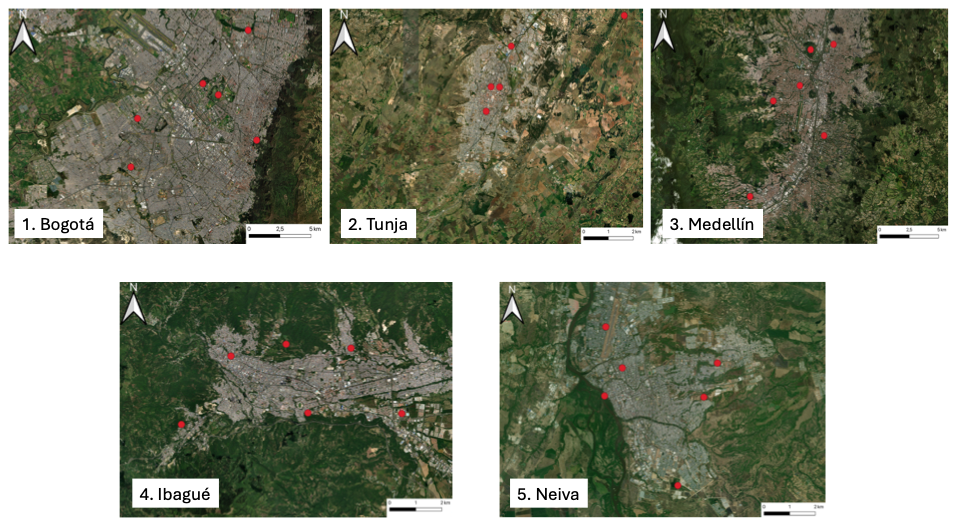

***Table S2.*** Percentage of impervious surface of each of the selected cities and study sites. Impervious surface for each selected city and study site was calculated using the Zonal Statistics tool in QGIS (QGIS Development Team 2018) on the 2024 Global Artificial Impervious Area (GAIA) raster covering the study region (Gong et al. 2020).

| City (Impervious surface %) | Site | Recreation allowed? | Impervious surface % |
| --- | --- | --- | --- |
| Bogotá (67. 5) | Humedal Burro | No | 9.49 |
|  | Humedal Córdoba | No | 1.86 |
|  | Simon Bolívar | Yes | 26.94 |
|  | Timiza | Yes | 25.79 |
|  | Javeriana | Yes | 60.54 |
|  | Libre | Yes | 42.6 |
| Medellín (65.55) | Cerro Volador | Yes | 1.88 |
|  | Jardín Botánico | No | 12.5 |
|  | Humedal Ditaires | No | 43.75 |
|  | Presidenta | Yes | 34.61 |
|  | Universidad de Medellín | Yes | 75 |
|  | UPB | Yes | 68.44 |
| Tunja (61.79) | Ecogranja | No | 1.82 |
|  | Reserva Malmo | No | 0.81 |
|  | Independencia | Yes | 42.8 |
|  | Muiscas | Yes | 47.23 |
|  | Pinzon | Yes | 57.89 |
|  | UPTC | Yes | 95.55 |
| Neiva (57.81) | Jardín Botánico | No | 0.25 |
|  | Mirador Mohan | No | 0.57 |
|  | Vergel | Yes | 25.56 |
|  | Cucaracha | Yes | 32.5 |
|  | UAN | Yes | 73.4 |
|  | Corhuila | Yes | 68.98 |
| Ibague (46.94 %) | San Jorge | No | 12.01% |
|  | Orquideas | No | 21.11% |
|  | Centenario | Yes | 36.21% |
|  | Deportivo | Yes | 38.32% |
|  | UniIbague | Yes | 65.49% |
|  | UTolima | Yes | 52.08% |

***Table S3.*** Number of labels (N_Samples) and, precision, recall and F1 of the model in detecting each of the species included in the developed CNN models. In total, 147 species of 16 orders, 34 families and 116 genera were included.

| Order | Family | Species | N_Samples | Precision | Recall | F1 |
| --- | --- | --- | --- | --- | --- | --- |
| Pelecaniformes | Ardeidae | Ardea alba | 1006 | 0.933 | 0.976 | 0.954 |
| Strigiformes | Strigidae | Asio clamator | 1027 | 0.988 | 0.988 | 0.988 |
| Anseriformes | Anatidae | Anser anser | 1038 | 0.968 | 0.977 | 0.972 |
| Strigiformes | Strigidae | Asio flammeus | 1052 | 0.936 | 0.971 | 0.953 |
| Psittaciformes | Psittacidae | Ara macao | 1003 | 0.963 | 0.981 | 0.972 |
| Gruiformes | Rallidae | Aramides cajaneus | 4993 | 0.958 | 0.839 | 0.894 |
| Psittaciformes | Psittacidae | Amazona ochrocephala | 39 | 0.991 | 0.959 | 0.975 |
| Anseriformes | Anatidae | Anas platyrhynchos | 1029 | 0.969 | 0.969 | 0.969 |
| Passeriformes | Tyrannidae | Atalotriccus pilaris | 1018 | 0.958 | 0.976 | 0.967 |
| Accipitriformes | Accipitridae | Accipiter striatus | 1012 | 0.994 | 0.896 | 0.943 |
| Apodiformes | Trochilidae | Amazilia tzacatl | 1185 | 0.967 | 0.962 | 0.964 |
| Passeriformes | Passerellidae | Arremon taciturnus | 1491 | 0.947 | 0.951 | 0.949 |
| Psittaciformes | Psittacidae | Amazona autumnalis | 1123 | 0.976 | 0.938 | 0.957 |
| Psittaciformes | Psittacidae | Ara severus | 1142 | 0.992 | 0.970 | 0.981 |
| Strigiformes | Strigidae | Asio stygius | 1071 | 0.994 | 0.967 | 0.980 |
| Accipitriformes | Accipitridae | Buteo albigula | 1010 | 0.428 | 0.870 | 0.574 |
| Passeriformes | Parulidae | Basileuterus culicivorus | 1010 | 0.389 | 0.760 | 0.515 |
| Passeriformes | Parulidae | Basileuterus tristriatus | 1042 | 0.395 | 0.332 | 0.361 |
| Psittaciformes | Psittacidae | Brotogeris jugularis | 1141 | 0.922 | 0.718 | 0.807 |
| Accipitriformes | Accipitridae | Buteo platypterus | 1023 | 0.956 | 0.675 | 0.792 |
| Passeriformes | Parulidae | Basileuterus rufifrons | 1500 | 0.985 | 0.398 | 0.567 |
| Passeriformes | Parulidae | Basileuterus tristriatus | 1044 | 0.597 | 0.672 | 0.632 |
| Cuculiformes | Cuculidae | Crotophaga ani | 1026 | 0.988 | 0.559 | 0.714 |
| Passeriformes | Corvidae | Cyanocorax affinis | 1010 | 0.992 | 0.272 | 0.427 |
| Apodiformes | Trochilidae | Colibri coruscans | 4377 | 0.822 | 0.306 | 0.446 |
| Passeriformes | Thraupidae | Coereba flaveola | 5444 | 0.728 | 0.721 | 0.724 |
| Passeriformes | Vireonidae | Cyclarhis gujanensis | 1193 | 0.971 | 0.893 | 0.930 |
| Passeriformes | Thraupidae | Catamenia homochroa | 1010 | 0.238 | 1.000 | 0.385 |
| Passeriformes | Icteridae | Chrysomus icterocephalus | 1374 | 0.912 | 0.966 | 0.938 |
| Anseriformes | Anatidae | Cairina moschata | 1013 | 0.962 | 0.918 | 0.940 |
| Apodiformes | Trochilidae | Coeligena coeligena | 171 | 0.983 | 0.787 | 0.874 |
| Piciformes | Picidae | Colaptes punctigula | 147 | 0.981 | 0.860 | 0.916 |
| Passeriformes | Turdidae | Catharus aurantiirostris | 1237 | 0.431 | 0.636 | 0.514 |
| Columbiformes | Columbidae | Columbina talpacoti | 1010 | 0.000 | 0.000 | 0.000 |
| Caprimulgiformes | Caprimulgidae | Chordeiles acutipennis | 1060 | 0.998 | 0.977 | 0.988 |
| Charadriiformes | Charadriidae | Charadrius vociferus | 1014 | 0.970 | 0.982 | 0.976 |
| Passeriformes | Corvidae | Cyanocorax yncas | 1059 | 0.969 | 0.984 | 0.976 |
| Anseriformes | Anatidae | Dendrocygna autumnalis | 1017 |  |  |  |
| Passeriformes | Thraupidae | Diglossa cyanea | 1028 | 0.971 | 0.990 | 0.980 |
| Passeriformes | Thraupidae | Diglossa humeralis | 1560 | 0.983 | 0.969 | 0.976 |
| Passeriformes | Thraupidae | Diglossa albilatera | 1058 | 0.908 | 0.866 | 0.887 |
| Passeriformes | Fringillidae | Euphonia chlorotica | 1017 | 0.973 | 0.947 | 0.959 |
| Passeriformes | Tyrannidae | Elaenia flavogaster | 1155 | 0.957 | 0.992 | 0.974 |
| Passeriformes | Tyrannidae | Elaenia frantzii | 1022 | 0.983 | 0.917 | 0.949 |
| Passeriformes | Tyrannidae | Euphonia laniirostris | 1202 | 0.934 | 0.940 | 0.937 |
| Accipitriformes | Accipitridae | Elanus leucurus | 1013 | 0.945 | 0.942 | 0.943 |
| Passeriformes | Fringillidae | Euphonia concinna | 1032 | 0.960 | 0.933 | 0.946 |
| Psittaciformes | Psittacidae | Eupsittula pertinax | 1016 | 1.000 | 0.960 | 0.980 |
| Passeriformes | Thraupidae | Eucometis penicillata | 1091 | 0.991 | 0.955 | 0.973 |
| Passeriformes | Tyrannidae | Empidonax virescens | 1682 | 0.942 | 0.942 | 0.942 |
| Gruiformes | Rallidae | Fulica americana | 92 | 0.875 | 0.919 | 0.896 |
| Psittaciformes | Psittacidae | Forpus conspicillatus | 1014 | 0.963 | 0.898 | 0.930 |
| Passeriformes | Thamnophilidae | Formicivora grisea | 1014 | 0.981 | 0.958 | 0.969 |
| Accipitriformes | Accipitridae | Geranoaetus albicaudatus | 1013 | 0.965 | 0.982 | 0.973 |
| Passeriformes | Grallariidae | Grallaricula ferrugineipectus | 1019 | 0.970 | 0.974 | 0.972 |
| Gruiformes | Rallidae | Gallinula galeata | 1575 | 0.972 | 0.936 | 0.954 |
| Strigiformes | Strigidae | Glaucidium jardinii | 1026 | 0.945 | 0.948 | 0.946 |
| Passeriformes | Grallariidae | Grallaricula nana | 71 | 0.949 | 0.973 | 0.961 |
| Passeriformes | Grallariidae | Grallaria ruficapilla | 1106 | 0.984 | 0.969 | 0.977 |
| Falconiformes | Falconidae | Herpetotheres cachinnans | 1017 | 0.852 | 0.894 | 0.872 |
| Passeriformes | Vireonidae | Hylophilus flavipes | 1506 | 0.992 | 0.982 | 0.987 |
| Passeriformes | Icteridae | Icterus chrysater | 1990 | 0.978 | 0.975 | 0.976 |
| Passeriformes | Tyrannidae | Legatus leucophaius | 1206 | 0.964 | 0.872 | 0.916 |
| Piciformes | Picidae | Leuconotopicus fumigatus | 1012 | 0.961 | 0.950 | 0.955 |
| Passeriformes | Furnariidae | Lepidocolaptes souleyetii | 1026 | 0.441 | 0.749 | 0.556 |
| Apodiformes | Trochilidae | Lesbia victoriae | 2791 | 0.984 | 0.953 | 0.968 |
| Passeriformes | Pipridae | Manacus manacus | 1064 | 0.630 | 0.650 | 0.640 |
| Passeriformes | Icteridae | Molothrus bonariensis | 1087 | 0.961 | 0.965 | 0.963 |
| Passeriformes | Tyrannidae | Myiozetetes cayanensis | 1165 | 0.267 | 0.948 | 0.417 |
| Strigiformes | Strigidae | Megascops choliba | 1131 | 0.983 | 0.929 | 0.956 |
| Passeriformes | Parulidae | Myiothlypis fulvicauda | 1098 | 0.965 | 0.987 | 0.976 |
| Piciformes | Picidae | Melanerpes formicivorus | 1010 | 0.481 | 0.956 | 0.640 |
| Passeriformes | Mimidae | Mimus gilvus | 1332 | 0.930 | 0.954 | 0.942 |
| Falconiformes | Falconidae | Milvago chimango | 1195 | 0.968 | 0.935 | 0.951 |
| Passeriformes | Thraupidae | Melanospiza bicolor | 1076 | 0.964 | 0.838 | 0.896 |
| Passeriformes | Parulidae | Myiothlypis fulvicauda | 1032 | 0.285 | 0.351 | 0.315 |
| Passeriformes | Tyrannidae | Mecocerculus leucophrys | 1027 | 0.513 | 0.942 | 0.664 |
| Passeriformes | Thamnophilidae | Myrmeciza longipes | 1022 | 0.960 | 0.958 | 0.959 |
| Passeriformes | Tyrannidae | Myiodynastes maculatus | 1070 | 0.978 | 0.977 | 0.977 |
| Passeriformes | Tyrannidae | Mionectes olivaceus | 1484 | 0.983 | 0.975 | 0.979 |
| Passeriformes | Tyrannidae | Myiotriccus ornatus | 10 | 0.282 | 0.893 | 0.428 |
| Passeriformes | Tyrannidae | Megarynchus pitangua | 1226 | 0.962 | 0.984 | 0.973 |
| Passeriformes | Tyrannidae | Machetornis rixosa | 1012 | 0.960 | 0.965 | 0.963 |
| Passeriformes | Turdidae | Myadestes ralloides | 1012 | 0.977 | 0.972 | 0.975 |
| Passeriformes | Tyrannidae | Myiozetetes similis | 1015 | 0.884 | 0.969 | 0.924 |
| Apodiformes | Trochilidae | Metallura tyrianthina | 1041 | 0.975 | 0.466 | 0.630 |
| Passeriformes | Tyrannidae | Myiarchus tuberculifer | 745 | 0.000 | 0.000 | 0.000 |
| Piciformes | Picidae | Melanerpes rubricapillus | 1086 | 0.978 | 0.929 | 0.953 |
| Caprimulgiformes | Caprimulgidae | Nyctidromus albicollis | 1455 | 0.932 | 0.932 | 0.932 |
| Tinamiformes | Tinamidae | Nothocercus julius | 1010 | 0.993 | 0.977 | 0.985 |
| Pelecaniformes | Ardeidae | Nycticorax nycticorax | 1030 | 0.472 | 0.284 | 0.355 |
| Galliformes | Cracidae | Ortalis columbiana | 1202 | 0.948 | 0.965 | 0.957 |
| Anseriformes | Anatidae | Oxyura jamaicensis | 1011 | 0.993 | 0.975 | 0.984 |
| Passeriformes | Hirundinidae | Orochelidon murina | 1320 | 0.939 | 0.979 | 0.958 |
| Passeriformes | Hirundinidae | Pygochelidon cyanoleuca | 1309 | 0.253 | 0.678 | 0.369 |
| Gruiformes | Rallidae | Porzana carolina | 1046 | 0.969 | 0.930 | 0.949 |
| Cuculiformes | Cuculidae | Piaya cayana | 1004 | 0.000 | 0.000 | 0.000 |
| Passeriformes | Troglodytidae | Pheugopedius fasciatoventris | 1113 | 0.955 | 0.925 | 0.939 |
| Apodiformes | Trochilidae | Phaethornis guy | 1001 | 0.963 | 0.936 | 0.949 |
| Gruiformes | Rallidae | Porphyrio martinica | 1010 | 0.000 | 0.000 | 0.000 |
| Psittaciformes | Psittacidae | Pionus menstruus | 1086 | 1.000 | 0.939 | 0.969 |
| Piciformes | Picidae | Picumnus olivaceus | 12 | 0.951 | 0.975 | 0.963 |
| Piciformes | Picidae | Picumnus olivaceus | 1026 | 0.492 | 0.927 | 0.643 |
| Passeriformes | Tyrannidae | Pitangus sulphuratus | 2336 | 0.499 | 0.932 | 0.650 |
| Passeriformes | Cardinalidae | Piranga rubra | 1213 | 0.937 | 0.695 | 0.798 |
| Passeriformes | Tyrannidae | Pyrocephalus rubinus | 1090 | 0.973 | 0.905 | 0.938 |
| Passeriformes | Icteridae | Quiscalus lugubris | 10 | 0.937 | 0.819 | 0.874 |
| Passeriformes | Thraupidae | Ramphocelus diamidatus | 1012 | 0.954 | 0.930 | 0.942 |
| Passeriformes | Thraupidae | Ramphocelus flammigerus | 1031 | 0.000 | 0.000 | 0.000 |
| Accipitriformes | Accipitridae | Rupornis magnirostris | 1130 | 0.981 | 0.910 | 0.945 |
| Passeriformes | Rhodinocichlidae | Rhodinocichla rosea | 1109 | 0.976 | 0.949 | 0.962 |
| Passeriformes | Parulidae | Setophaga castanea | 1043 | 0.961 | 0.988 | 0.974 |
| Passeriformes | Parulidae | Setophaga cerulean | 1012 | 0.966 | 0.966 | 0.966 |
| Passeriformes | Parulidae | Setophaga petechia | 1311 | 0.882 | 0.934 | 0.907 |
| Passeriformes | Thraupidae | Sicalis flaveola | 4866 | 0.987 | 0.920 | 0.952 |
| Passeriformes | Thraupidae | Stilpnia heinei | 1050 | 0.760 | 0.586 | 0.662 |
| Caprimulgiformes | Caprimulgidae | Systellura longirostris | 1031 | 0.905 | 0.903 | 0.904 |
| Passeriformes | Icteridae | Sturnella magna | 662 | 0.997 | 0.950 | 0.973 |
| Passeriformes | Thraupidae | Saltator maximus | 1568 | 0.956 | 0.949 | 0.953 |
| Passeriformes | Furnariidae | Synallaxis brachyura | 1041 | 0.805 | 0.919 | 0.858 |
| Passeriformes | Thraupidae | Saltator olivascens | 1166 | 0.952 | 0.943 | 0.947 |
| Passeriformes | Fringillidae | Spinus psaltria | 1486 | 0.905 | 0.784 | 0.840 |
| Passeriformes | Fringillidae | Spinus spinescens | 1361 | 0.953 | 0.942 | 0.947 |
| Passeriformes | Thraupidae | Saltator striatipectus | 1409 | 0.876 | 0.963 | 0.917 |
| Passeriformes | Parulidae | Setophaga striata | 1010 | 0.945 | 0.992 | 0.968 |
| Passeriformes | Thraupidae | Sicalis luteola | 1023 | 0.969 | 0.919 | 0.943 |
| Apodiformes | Apodidae | Streptoprocne zonaris | 1116 | 0.428 | 0.626 | 0.509 |
| Passeriformes | Troglodytidae | Troglodytes aedon | 2594 | 0.956 | 0.920 | 0.938 |
| Passeriformes | Tyrannidae | Todirostrum cinereum | 199 | 0.893 | 0.789 | 0.838 |
| Passeriformes | Thraupidae | Thraupis episcopus | 5090 | 0.845 | 0.816 | 0.830 |
| Passeriformes | Turdidae | Turdus fuscater | 7544 | 0.967 | 0.921 | 0.943 |
| Passeriformes | Turdidae | Turdus grayi | 745 | 0.000 | 0.000 | 0.000 |
| Passeriformes | Thraupidae | Tangara gyrola | 1016 | 0.961 | 0.976 | 0.969 |
| Passeriformes | Turdidae | Turdus ignobilis | 1222 | 0.951 | 0.909 | 0.929 |
| Passeriformes | Turdidae | Turdus leucomelas | 2455 | 0.990 | 0.849 | 0.914 |
| Passeriformes | Tyrannidae | Tyrannus melancholicus | 2551 | 0.986 | 0.931 | 0.958 |
| Passeriformes | Thamnophilidae | Thamnophilus multistriatus | 1220 | 0.785 | 0.528 | 0.631 |
| Passeriformes | Thraupidae | Tiaris olivaceus | 1013 | 0.959 | 0.877 | 0.916 |
| Passeriformes | Thraupidae | Thraupis palmarum | 1880 | 0.984 | 0.963 | 0.973 |
| Passeriformes | Tyrannidae | Tolmomyias poliocephalus | 1053 | 0.983 | 0.960 | 0.972 |
| Passeriformes | Tyrannidae | Tyrannus savana | 1083 | 0.995 | 0.987 | 0.991 |
| Passeriformes | Thraupidae | Tersina viridis | 1013 | 0.000 | 0.000 | 0.000 |
| Charadriiformes | Charadriidae | Vanellus chilensis | 1353 | 0.987 | 0.981 | 0.984 |
| Passeriformes | Vireonidae | Vireolanius eximius | 1015 | 0.000 | 0.000 | 0.000 |
| Passeriformes | Vireonidae | Vireo olivaceus | 1027 | 0.901 | 0.900 | 0.900 |
| Passeriformes | Passerellidae | Zonotrichia capensis | 4493 | 0.978 | 0.971 | 0.974 |
| Passeriformes | Tyrannidae | Zimmerius chrysops | 1024 | 0 | 0 | 0 |

***Table S4.*** Estimates, Standard errors (SE), degrees of freedom and P-values from pairwise comparisons between the different model configurations evaluated in this study. Bold values indicate statistical significance at α=0.05. N=10.584.

| Comparison | Estimate | SE | DF | T-Ratio | P.value |
| --- | --- | --- | --- | --- | --- |
| Buffer05 - Buffer1 | 0.185 | 0.043 | 10422.852 | 4.3 | 0.0004588 |
| Buffer05 - LogMel | -1.87 | 0.042 | 10422.852 | -44.072 | <0.01 |
| buffer05 - Non_Augmentated | -0.567 | 0.064 | 10422.851 | -8.854 | <0.01 |
| buffer05 - pcen | -0.778 | 0.043 | 10422.851 | -18.279 | <0.01 |
| buffer05 - presence | -1.094 | 0.042 | 10422.851 | -25.791 | <0.01 |
| buffer05 - single | 0.729 | 0.0423 | 10422.851 | 17.216 | <0.01 |
| buffer05 - weighted | -0.679 | 0.042 | 10422.851 | -15.955 | <0.01 |
| buffer1 - LogMel | -2.054 | 0.0424 | 10422.851 | -48.433 | <0.01 |
| buffer1 - Non_Augmentated | -0.753 | 0.064 | 10422.851 | -11.742 | <0.01 |
| buffer1 - pcen | -0.963 | 0.046 | 10422.851 | -22.627 | <0.01 |
| buffer1 - presence | -1.279 | 0.042 | 10422.851 | -30.155 | <0.01 |
| buffer1 - single | 0.544 | 0.0423 | 10422.851 | 12.847 | <0.01 |
| buffer1 - weighted | -0.865 | 0.0425 | 10422.851 | -20.3 | <0.01 |
| LogMel - Non_Augmentated | 1.302 | 0.063 | 10422.851 | 20.464 | <0.01 |
| LogMel - pcen | 1.092 | 0.042 | 10422.851 | 26.045 | <0.01 |
| LogMel - presence | 0.775 | 0.0417 | 10422.851 | 18.578 | <0.01 |
| LogMel - single | 2.59 | 0.041 | 10422.851 | 62.346 | <0.01 |
| LogMel - weighted | 1.189 | 0.041 | 10422.851 | 28.366 | <0.01 |
| Non_Augmentated - pcen | -0.21 | 0.063 | 10422.851 | -3.294 | 0.022 |
| Non_Augmentated - presence | -0.525 | 0.063 | 10422.851 | -8.25 | <0.01 |
| Non_Augmentated - single | 1.297 | 0.06 | 10422.851 | 21.288 | <0.01 |
| Non_Augmentated - weighted | -0.111 | 0.063 | 10422.851 | -1.752 | 0.652 |
| pcen - presence | -0.315 | 0.041 | 10422.851 | -7.538 | <0.01 |
| pcen - single | 1.507 | 0.041 | 10422.851 | 36.007 | <0.01 |
| pcen - weighted | 0.098 | 0.042 | 10422.851 | 2.336 | 0.274 |
| presence - single | 1.823 | 0.04 | 10422.851 | 43.71 | <0.01 |
| presence - weighted | 0.414 | 0.041 | 10422.851 | 9.876 | <0.01 |
| single - weighted | -1.409 | 0.041 | 10422.851 | -33.629 | <0.01 |

***Table S5.*** Comparison between the performance of our CNN and BirdNET (Kahl et al., 2021) in identifying species in tropical urban-rural gradients. The table contrasts performance in terms of Micro-F1, Macro-F1, and the number of predicted positives. Micro-F1 reflects overall classification performance weighted by class frequency, whereas Macro-F1 represents the unweighted mean performance across classes, emphasizing performance on less frequent species.

|  |  | Micro-F1 | | | Macro-F1 | | |
| --- | --- | --- | --- | --- | --- | --- | --- |
| Model | Predicted positives | Precision | Recall | F1 | Precision | Recall | F1 |
| CNN 3 + ConveNeXt | 3904 | 0.853 | 0.740 | 0.792 | 0.651 | 0.471 | 0.547 |
| BirdNET | 775 | 0.754 | 0.077 | 0.140 | 0.260 | 0.086 | 0.114 |

***Table S6.*** Species uniquely detected by each method. In total, 55 species were uniquely identified by the CNN and 43 by the Point Counts.

| Species | Method |
| --- | --- |
| Columba livia | PointCount |
| Zenaida auriculata | PointCount |
| Molothrus bonariensis | PointCount |
| Columbina talpacoti | PointCount |
| Phimosus infuscatus | PointCount |
| Tachyphonus rufus | PointCount |
| Momotus aequatorialis | PointCount |
| Thraupis palmarium | PointCount |
| Aramides cajaneus | PointCount |
| Ramphocelus dimidiatus | PointCount |
| Thamnophilus doliatus | PointCount |
| Basileuterus delattrii | PointCount |
| Coragyps atratus | PointCount |
| Ardea ibis | PointCount |
| Piranga rubra | PointCount |
| Sicalis citrina | PointCount |
| Camptostoma obsoletum | PointCount |
| Sayornis nigricans | PointCount |
| Chrysuronia goudoti | PointCount |
| Euphonia concinna | PointCount |
| Tolmomyias sulphurescens | PointCount |
| Dendrocygna autumnalis | PointCount |
| Butorides striata | PointCount |
| Tyrannus sp | PointCount |
| Tiaris olivaceus | PointCount |
| Myiozetetes similis | PointCount |
| Saltator maximus | PointCount |
| Saucerottia saucerottei | PointCount |
| Catharus aurantiirostris | PointCount |
| Synallaxis albescens | PointCount |
| Catharus ustulatus | PointCount |
| Pyrocephalus obscurus | PointCount |
| Tangara vassorii | PointCount |
| Chlorospingus flavopectus | PointCount |
| Vireo olivaceus | PointCount |
| Vireo flavoviridis | PointCount |
| Quiscalus mexicanus | PointCount |
| Mniotilta varia | PointCount |
| Oxyura jamaicensis | PointCount |
| Ocreatus underwoodii | PointCount |
| Diglossa cynea | PointCount |
| Zimmerius chrysops | PointCount |
| Eupsittula pertinax | CNN |
| Asio clamator | CNN |
| Basileuterus culicivorus | CNN |
| Megascops choliba | CNN |
| Systellura longirostris | CNN |
| Pyrocephalus rubinus | CNN |
| Anser anser | CNN |
| Cyanocorax yncas | CNN |
| Nycticorax nycticorax | CNN |
| Cyanocorax affinis | CNN |
| Thraupis palmarum | CNN |
| Nyctidromus albicollis | CNN |
| Mionectes olivaceus | CNN |
| Catharus aurantiirostris | CNN |
| Grallaria ruficapilla | CNN |
| Tersina viridis | CNN |
| Metallura tyrianthina | CNN |
| Thamnophilus multistriatus | CNN |
| Buteo albigula | CNN |
| Turdus ignobilis | CNN |
| Herpetotheres cachinnans | CNN |
| Saltator striatipectus | CNN |
| Saltator olivascens | CNN |
| Ara severus | CNN |
| Basileuterus rufifrons | CNN |
| Amazona automnalis | CNN |
| Spinus spinescens | CNN |
| Euphonia chlorotica | CNN |
| Amazona ochrocephala | CNN |
| Quiscalus lugubris | CNN |
| Hylophilus flavipes | CNN |
| Myiodynastes maculatus | CNN |
| Asio stygius | CNN |
| Ardea alba | CNN |
| Ara macao | CNN |
| Eucometis penicillata | CNN |
| Arremon taciturnus | CNN |
| Porphyrio martinica | CNN |
| Piaya cayana | CNN |
| Aramides cajenus | CNN |
| Myrmeciza longipes | CNN |
| Pheugopedius fasciatoventris | CNN |
| Formicivora grisea | CNN |
| Asio flammeus | CNN |
| Myadestes ralloides | CNN |
| Setophaga striata | CNN |
| Lesbia victoriae | CNN |
| Euphonia laniirostris | CNN |
| Phaethornis guy | CNN |
| Elaenia frantzii | CNN |
| Setophaga cerulean | CNN |
| Ramphocelus flammigerus | CNN |
| Buteo platypterus | CNN |
| Myiotriccus ornatus | CNN |
| Mecocerculus leucophrys | CNN |
